# Early-Life Gut Inflammation Drives Sex-Dependent Shifts in the Microbiome-Endocrine-Brain Axis

**DOI:** 10.1101/2024.05.23.595618

**Authors:** Olivia Sullivan, Claire Sie, Katharine M. Ng, Sophie Cotton, Cal Rosete, Jordan E. Hamden, Ajay Paul Singh, Kristen Lee, Jatin Choudhary, Jennifer Kim, Huaxu Yu, Charlotte A. Clayton, Natalia A. Carranza Garcia, Kateryna Voznyuk, Brian D. Deng, Nadine Plett, Sana Arora, Hans Ghezzi, Tao Huan, Kiran K. Soma, John-Paul J. Yu, Carolina Tropini, Annie Vogel Ciernia

## Abstract

Despite recent advances in understanding the connection between the gut microbiota and the adult brain, there remains a wide knowledge gap in how gut inflammation impacts brain development. We hypothesized that intestinal inflammation in early life would negatively affect neurodevelopment through dysregulation of microbiota communication to the brain. We therefore developed a novel pediatric chemical model of inflammatory bowel disease (IBD), an incurable condition affecting millions of people worldwide. IBD is characterized by chronic intestinal inflammation, and has comorbid symptoms of anxiety, depression and cognitive impairment. Significantly, 25% of patients with IBD are diagnosed during childhood, and the effect of chronic inflammation during this critical period of development is largely unknown. This study investigated the effects of early-life gut inflammation induced by DSS (dextran sulfate sodium) on a range of microbiota, endocrine, and behavioral outcomes, focusing on sex-specific impacts. DSS-treated mice exhibited increased intestinal inflammation, altered microbiota membership, and changes in microbiota-mediated circulating metabolites. The majority of behavioral measures were unaffected, with the exception of impaired mate-seeking behaviors in DSS-treated males. DSS-treated males also showed significantly smaller seminal vesicles, lower circulating androgens, and decreased intestinal hormone-activating enzyme activity. In the brain, microglia morphology was chronically altered with DSS treatment in a sex-specific manner. The results suggest that early-life gut inflammation causes changes in gut microbiota composition, affecting short-chain fatty acid (SCFA) producers and glucuronidase (GUS) activity, correlating with altered SCFA and androgen levels. The findings emphasize the developmental sensitivity to inflammation-induced changes in endocrine signalling and underscore long-lasting physiological and microbiome changes associated with juvenile IBD.

**Highlights:** Early-life gut inflammation produces sex-specific effects on i) microbiome, ii) sex hormones and iii) behaviour.

Both sexes show disrupted gut bacterial members that regulate sex hormone levels.

Male mice demonstrate deficits in mate seeking, which may be mediated by reduced androgen levels.

Both male and female mice demonstrate shifts in hippocampal microglial morphology.

## Introduction

Inflammatory bowel disease (IBD) is a group of chronic gastrointestinal (GI) tract conditions that affects over 6.8 million people globally and remains incurable (Alatab et al., 2020). The risk of IBD is correlated with industrialization and, as such, is globally rising in prevalence (Kaplan and Windsor, 2021). Genetic factors account for less than one-third of IBD risk, underscoring the importance of environmental factors such as gut microbiota composition, dietary habits, smoking, and stress levels on disease development (Ananthakrishnan, 2013; Ye et al., 2015). Individuals with IBD frequently undergo cycles of intense inflammatory flare-ups and remission periods (Roda et al., 2020). Over 25% of patients with IBD are diagnosed during childhood or adolescence with symptoms that persist through adulthood (Däbritz et al., 2017). Importantly, little is known about the specific impacts of pediatric-onset IBD on the developing brain. Children and adolescents with IBD have an increased risk of developing anxiety and depression, experience a lower quality of life, and face educational and social challenges along with mild cognitive impairments (Attree et al., 2003; Mackner et al., 2013; Thavamani et al., 2019). Furthermore, neuroimaging studies in adult female and male patients with IBD have revealed altered brain structure and function (Bao et al., 2015; Lv et al., 2017) some of which correlate with neuropsychiatric impairments (Hou et al., 2020), highlighting the need for similar research in pediatric populations.

Research into inflammatory phenotypes through various animal models – including genetic, chemically induced, and adoptive transfer T cell models – has significantly advanced our understanding of IBD (Wirtz and Neurath, 2007). The widely-used dextran sodium sulfate (DSS)-induced colitis model damages the gut epithelial lining and mimics the gut inflammation and microbiota changes seen in human ulcerative colitis (UC), one of the two main types of IBD (Chassaing et al., 2014). Adult mice treated with DSS have increased anxiety (Nyuyki et al., 2018) and reduced sucrose preference, indicative of a depression endophenotype (Zhou et al., 2023). Additionally, active gut inflammation in adult mice led to anxiety-like behavior and recognition memory impairments (Emge et al., 2016) and social impairments (Brown et al., 2024). Importantly, anxiety and memory deficits were reversed upon resolution of acute inflammation (Emge et al., 2016), which is encouraging for therapeutic resolution of neurophysiological symptons in humans with IBD. While DSS-colitis studies predominantly focus on adult rodents and capture many of the extraintestinal manifestations of IBD seen in adult human patients, research in juvenile rodent models remains limited. In contrast to adult mice, a recent DSS-colitis model in juvenile mice revealed persistent cognitive deficits and anxiety-like behaviors even upon GI-symptom resolution (Salvo et al., 2020). These observations suggest that GI inflammation during developmental periods may have unique and long-lasting effects on brain development, leading to distinct behavioral impacts in adulthood.

How GI inflammation triggers brain and behavior dysfunction is not well understood, but is theorized to involve altered signaling along the gut-brain-axis. The gut microbiota can signal to the brain by immune cell trafficking, peripheral nerve activity, and release of metabolites that circulate to the brain (Banfi et al., 2021; Günther et al., 2021). One of the key brain cells responsive to gut microbe derived metabolites are microglia, the brain’s resident innate immune cells. Microglia sculpt synaptic physiology during brain development (Sullivan and Ciernia, 2022), are acutely sensitive to changes in microbiota in early life (Erny et al., 2015; Thion et al., 2017) and are chronically functionally altered from early life inflammation (Vogel Ciernia et al., 2018). Microbiota-derived short chain fatty acids (SCFAs) can enter the blood stream and cross the blood brain barrier to impact microglia development and function (Cryan and Dinan, 2012; Erny et al., 2015; Sullivan and Ciernia, 2022). In both adult and juvenile IBD mouse models, microglia adopt a pro-inflammatory phenotype characterized by altered cellular morphology and increased pro-inflammatory cytokine secretion (Caetano-Silva et al., 2024; Gampierakis et al., 2021; Salvo et al., 2020; Sroor et al., 2019; Talley et al., 2021). Together, this suggests that early life changes to microbiota-derived signalling to brain microglia may negatively impact brain development in IBD.

Beyond impacting brain function and behavior, pediatric IBD poses several unique challenges in other aspects of development, including delayed onset of puberty in both males and females (Ballinger et al., 2003). While undernutrition was inititally thought to be the main driver of delayed puberty, restoring nutritional status fails to normalize the timing of puberty onset, suggesting the involvement of additional factors (Azooz et al., 2001; Brain and Savage, 1994; DeBoer et al., 2010; Deboer and Li, 2011). Animal models of IBD have linked intestinal inflammation to disruption in testosterone levels, estrogen receptor function, and reproductive organ size, with effects that extend beyond what is seen with undernutrition alone (Azooz et al., 2001; Ballinger et al., 2003; DeBoer et al., 2010; Deboer and Li, 2011). Similarly, findings in human studies show that some patients with IBD (Klaus et al., 2008; Tigas and Tsatsoulis, 2012) and irritable bowel syndrome (Rastelli et al., 2022) have disrupted sex hormone levels and higher rates of sexual dysfunction (Chen et al., 2024; Gaidos et al., 2020). One connection between inflammation and altered hormone levels is thought to be mediated by the gut microbiota, which acts as a regulator of estrogen and androgen synthesis and metabolism (Colldén et al., 2019; He et al., 2021; Maffei et al., 2022; Neuman et al., 2015). In mouse models of IBD, the loss of bacterial species such as *Faecalibacterium prausnitzii* (Sokol et al., 2008) and *Roseburia hominis*, known for producing anti-inflammatory SCFAs, is associated with changes in sex hormone levels (Sui et al., 2021; Torres et al., 2018). The depletion of specific bacteria such as *F. prausnitzii* in IBD, which are crucial for converting estrogen to its active form, can directly impact hormone availability, potentially affecting the onset and progression of sexual development (Ervin et al., 2019). Additionally, the recent discovery of testosterone-degrading bacteria such as *Mycobacterium* contributing to inflammation in IBD underscores the microbiota’s profound impact on sexual development, suggesting avenues for targeted microbial interventions to mitigate impairments in IBD (Li et al., 2022; Naser et al., 2014). Altogether, these studies highlight the complex relationships between the gut microbiota and sexual development (Lavelle and Sokol, 2020; Machiels et al., 2014).

There is a critical gap in understanding the interactions between early life intestinal inflammation, brain function, and reproductive development. To bridge this gap, here we introduce a mouse model of pediatric-onset IBD that captures recurrent episodes of gut inflammation throughout childhood and adolescence, modeling the human condition. In both sexes, we assessed the acute and chronic effects of early-life gut inflammation on gut microbiota composition, short-chain fatty acid production, a wide variety of behaviors, sex steroid levels, brain white matter tract integrity and microglial morphology, offering a comprehensive view of IBD’s impact on juvenile development. Our findings reveal novel sex-specific changes in reproductive behavior, alterations in microbiota metabolites and hormone signaling, and microglial morphology adaptations. These results highlight the unique long-term consequences for human patients with early-onset IBD, emphasizing a need for early diagnosis and microbiome-targeted intervention strategies.

## Methods

### Animals

Experimentally naïve 3-week-old C57BL/6J wildtype mice (strain #000664, Jackson Laboratories) were housed in groups of 3-5 with sex and treatment-matched cage mates. Mice were housed in ventilated cages under a 12:12 light-dark cycle with lights on at 0700h and lights off at 1900h in a temperature (22°C) and humidity (40-60%) controlled environment. Purina LabDiet 5k67 diet and reverse osmosis chlorinated water were given *ad libitum* throughout the experiment unless stated otherwise for specific tasks. All housing conditions and testing procedures were in accordance with the guidelines of the Canadian Council on Animal Care, and all protocols were approved by the Animal Care Committee of the University of British Columbia *(protocols A19-0078, A23-0115, and A23-0086)*.

### Pediatric IBD mouse model

Mice randomly assigned to the treatment groups were administered three rounds of 5-day treatments of 3-3.5% colitis-grade DSS (36-50 kDa, MP Biomedicals #160110) while the vehicle control group (VEH) received water only. Due to the reported batch-dependent effects of DSS, the optimal concentration of DSS to cause colitis was determined in preceding pilot experiments (Eichele and Kharbanda, 2017). The treatment periods occurred during postnatal days (P) P21-26, P42-47, and P63-68 (Figure 1A). During treatment and for three days following each treatment, mice were weighed daily to monitor for weight loss (Supplemental Table 2). Regular monitoring of weight was continued during the recovery period intervals. One cohort of mice was euthanized three days after the final DSS treatment on P71 (n=4-5/treatment/sex). The remaining two cohorts of mice underwent behavioral testing as young adults between P85-P130 (n=11-13/treatment/sex).

**Figure 1.**
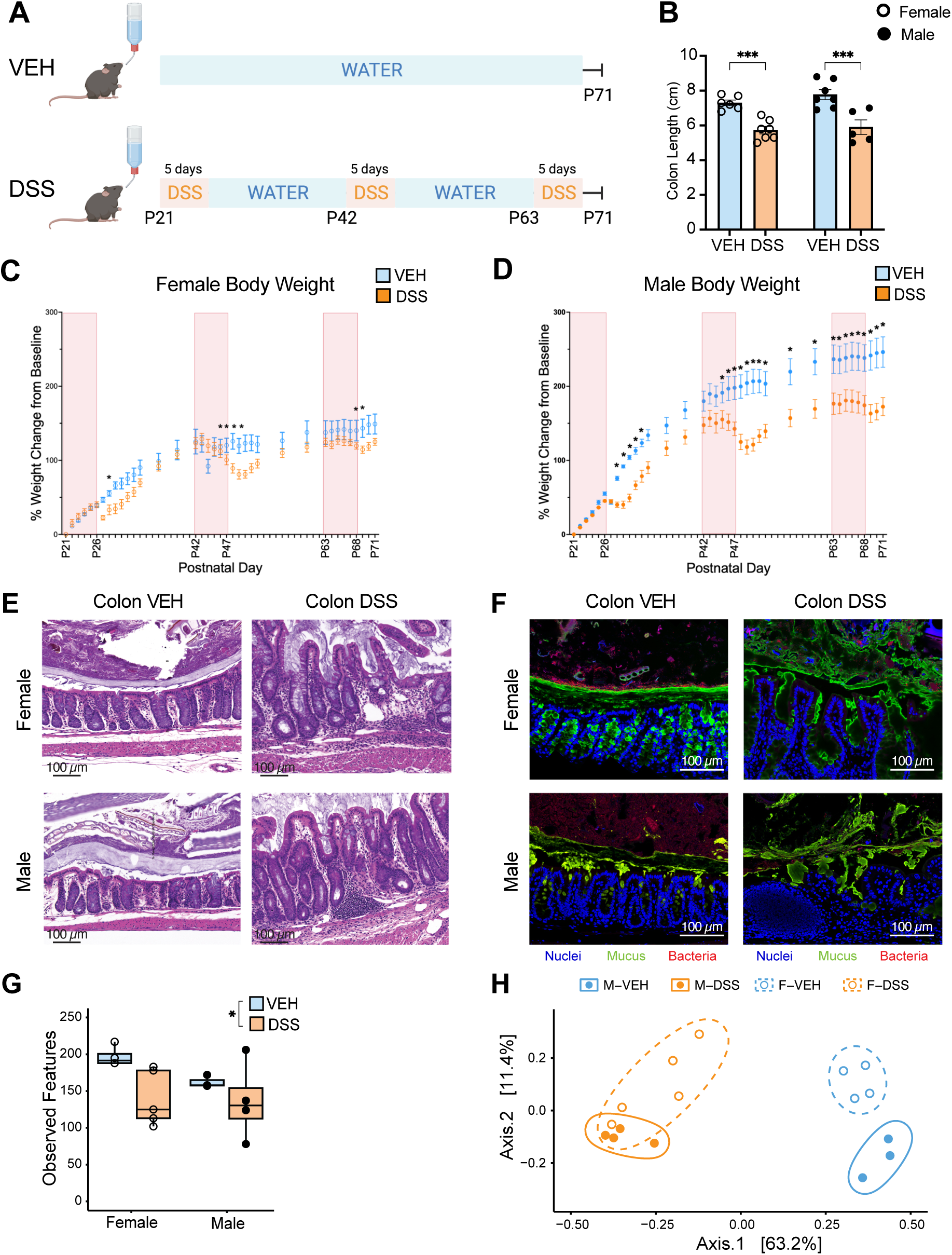
Juvenile mouse DSS treatment models early-life intestinal inflammation. *A*. Experimental schematic. Control mice were given water throughout (VEH), while treatment mice were given 3 rounds of 5-day long DSS treatments beginning at P21, P42, and P63 (DSS). *B.* Post-treatment colon lengths measured at P71, three days after the final round of DSS, were shorter in DSS-treated mice. Each graph bar and error bar represent the mean ± SEM, n=5-7/group. *C, D.* Female and male percent weight change from baseline (P21) starting weight. Each point and error bar represents the mean ± SEM, n=11-13/group. *E.* Representative images of hematoxylin and eosin-stained colonic tissues from mice collected at P71 confirm that DSS treatment induced intestinal inflammation. Scale bar, 100 µm. *F.* Representative fluorescence *in situ* hybridization (FISH) images from VEH and DSS treated mice shows gut disruption in DSS treated mice. Scale bar, 100 µm. Colors: host tissue (DAPI, blue), mucus (UEA-1, green), and bacteria (Eub-338, red). *G.* Microbiota alpha diversity as measured by number of observed unique features (a proxy for number of species) per sample is significantly lower in DSS-treated mice, n=3-5/group. *H.* Principal Component Analysis of Bray-Curtis diversity index shows separation by DSS treatment, n=3-5/group. \**p*<.05. \*\*\**p*<.0001.

### Serum and organ collection

Animals were deeply anesthetized with 5% isoflurane, and blood was collected by cardiac puncture. Whole blood was collected into BD Microtainer^TM^ Serum Separator Tubes (BD) and allowed to coagulate for at least 30 minutes at room temperature according to manufacturer instructions. To isolate serum for later analyses, blood samples were spun down at 1200 x g for 20 minutes at 4°C, and the serum was removed and placed in a new tube for storage at −80°C. Animals were then transcardially perfused with 1X phosphate buffered saline (PBS). Intestinal measures were conducted as follows: first, intestines were dissected out from the gastroduodenal junction to the rectum. Colon length was then measured from the ileocecal junction to the rectum. The contents of the cecum were gently scraped and collected into a sterile tube, then immediately flash-frozen on dry ice. The colon was manipulated into rolls in histological cassettes and immediately fixed in freshly prepared Methacarn solution (60% dry methanol, 30% chloroform, 10% glacial acetic acid) (Ng and Tropini, 2021). Brains were collected and drop fixed in 4% paraformaldehyde (PFA) for 24 hours at 4°C and then transferred to 30% sucrose solution and kept at 4°C until sectioned. For brain connectivity analysis, mice were perfused with PBS followed by 4% PFA and brains were dissected out and then stored in 4% PFA until scanning.

### Gut staining and histological scoring

Within 2 weeks of fixation in Methacarn solution, intestinal tissues were processed and infiltrated with paraffin wax as previously described (Ng and Tropini, 2021). Briefly, tissues were washed twice with absolute methanol for 30 minutes each, followed by two washes with absolute ethanol for 20 minutes each. Next, tissues were washed twice with xylene for 15 minutes before being placed in melted paraffin wax and incubated for 2 hours at 60°C. Samples were then sent to the BC Children’s Hospital Histology Core for embedding in paraffin wax and sectioning into 4 – 5 µm sections. Histological staining was performed with hematoxylin and eosin as previously described (Stahl et al., 2017). Finally, slides were imaged using a PANNORAMIC® MIDI II (3DHISTECH).

### Mucus and bacteria visualization

Visualization of gut microbes was performed as previously described (Ng and Tropini, 2021). Briefly, to deparaffinize tissues, slides were heated at 60°C in Coplin jars for 10 minutes, followed by incubation in xylene that was pre-warmed to 60°C. A second xylene wash was performed at room temperature for 10 minutes, then slides were placed in 99.5% ethanol for 5 minutes. After allowing slides to air-dry, tissues were encircled using a liquid blocker PAP pen. FISH probe hybridization solution (20 mM Tris-HCl, pH 7.4; 0.9 M NaCl; 0.1% sodium dodecylsulfate in nuclease-free water) was pre-warmed to 50°C before addition of Eub-388 (5’-AF488-GCTGCCTCCCGTAGGAGT-3’) to a final concentration of 10 ng/µL (Amann et al., 1990). Slides were then incubated for 3 hours at 50°C in a humid chamber. Following incubation, tissues were washed once with FISH washing buffer (20 mM Tris-HCl, pH 7.4; 0.9 M NaCl, pre-warmed to 50°C) for 10 minutes at 50°C, then additionally with 1x PBS. DNA and intestinal mucus were counter-stained using 10 µg/mL DAPI and 40 µg/mL UEA-1-Rhodamine Red respectively in PBS for 45 minutes at 4°C. Finally, slides were washed thrice with PBS before mounting with ProLong Gold AntiFade mounting medium (Invitrogen). Fluorescence images were captured using a Zeiss LSM900 confocal microscope using a 100x objective lens.

### Microbiome sequencing and analysis

Microbiome composition was determined via sequencing of the 16S rRNA hypervariable region. Total DNA was extracted from 30-60 mg of cecal contents using the DNeasy PowerSoil Pro kit (Qiagen) in accordance with manufacturer protocols. Quantification of extracted DNA concentration prior to library preparation was performed using the Quant-iT^TM^ dsDNA HS Assay (Thermo Fisher). 16S library preparation was conducted at either the Gut4Health facility at BC Children’s Hospital or the Biofactorial High-Throughput Biology (Bio!) Facility at the University of British Columbia. For samples collected 3 days after cessation of DSS, library preparation targeting the V4 hypervariable region of the 16S rRNA gene was performed at the Gut4Health facility using established protocols (De Wolfe and Wright, 2023). Amplification of the V4 region was performed with 515F/926R primers (515F, 5’-GTGYCAGCMGCCGCGGTAA-3’; 926R, 5’-CCGYCAATTYMTTTRAGTTT-3’). Pooled libraries were then submitted to the Bio! facility, where sequencing was performed on the Illumina MiSeq^TM^ platform with v2 2 x 250 bp paired-end read chemistry. For all other samples, library preparation targeting the V4-V5 16S rRNA genes was performed at the Bio! Facility as previously described (Ng et al., 2023), and sequencing was run on the Illumina MiSeq^TM^ platform with v3 2 x 300 bp paired-end read chemistry. FASTQC (Andrews, 2010) was run on the generated FASTQ files to assess read quality. Reads were then imported into QIIME2-2023.2 for subsequent analyses (Bolyen et al., 2019). Denoising and quality filtering was performed using DADA2 (via q2-dada2), and reads were trimmed to remove primer sequences while maintaining mean Phred quality scores >Q30 (Callahan et al., 2016). Multiple sequence alignment and phylogenetic tree generation was performed using MAFFT (via q2-alignment) and FastTree2 respectively (via q2-phylogeny) (Katoh et al., 2002; Price et al., 2010). Using the QIIME classification plugin (q2-feature-classifier), amplicon sequence variants (ASVs) were classified via a naïve Bayes machine-learning taxonomic classifier against the SILVA 138 99% identity reference sequence database (Quast et al., 2013).

Further analyses were conducted using R v4.2.2 (R Core team, 2021). Analyses were visualized through the tidyverse (Wickham et al., 2019) and ggplot2 (Hadley, 2016) packages. For sample rarefaction, calculation, and visualization of alpha and beta diversity metrics, the phyloseq (McMurdie and Holmes, 2013), ggpubr (Kassambara, 2023), and vegan (Dixon, 2003) packages were employed. Differential abundance analysis (DAA) was conducted using the metagenomeseq package (Paulson et al., 2013) and Analysis of Composition of Microbiomes (ANCOM) (Mandal et al., 2015) packages through the microbiomeMarker package (Cao et al., 2022). The relative abundance of those taxa identified as significantly differentially abundant were then plotted with FDR-corrected p-values obtained with metagenomeseq. Differentially abundant taxa were plotted using the ComplexHeatmap (Gu et al., 2016) package. The relative abundance of select differentially abundant taxa was then plotted against serum hormone and SCFA concentrations, and Spearman’s correlation was calculated using smplot2 (Min and Zhou, 2021).

### Behavior testing

All behavior testing was performed under white light (600 lux) between the hours of 0900-1800 except for the sucrose preference test, which is a 24-hour task. Animals were habituated to the researchers after the anxiety tasks but prior to all other tests through 3 days of handling (1-2 min/mouse). For each behavior task, animals were first habituated to room conditions for 30 minutes before testing. For all tasks that did not involve bedding, the apparatus was cleaned with diluted saber solution (Vert-2-Go saber diluted 1:16 in water) between mice and between stages. ANY-maze tracking software (Stoelting) was used to record videos and track animal movement during the task. Mice began testing 17 days following the final round of DSS with the exception of a subset cohort, which underwent only the urine preference test and mating behavior test beginning 28 days following the final round of DSS treatment.

#### Urine Preference Test (UPT)

Mice were placed in the centre of an open-top box (40 cm x 40 cm x 40 cm, L x W x H) containing two 10 cm diameter inverted pencil cups in opposite corners. A petri dish with blotting paper (1 cm^2^) was placed under each pencil cup. Mice were given a 10-minute habituation period where 20 µL of deionized water was dropped onto both blotting papers. Following habituation, mice were given another 10-minutes to explore the two cups, this time with 20 µL of pooled male or female C57BL/6 mouse urine dropped onto one of the blotting papers and 20 µL of deionized water dropped onto the other. In the final stage, mice were given another 10-minutes to explore the two cups, with the urine cup location and urine sex switching from the previous stage (Malkesman et al., 2010). Presentation order of same or opposite sex urine as well as location of urine in the first and second stages was counterbalanced across mice and treatment conditions. ANY-maze tracked the total investigation time when the test mouse’s nose was within 1 cm of the inverted pencil cup. Exclusion criteria for data collected during the urine preference test include side preference during habituation (defined as a discrimination index (DI) over 20 or under −20, where DI is the difference in time spent investigating the stimulus cups divided by the total time spent investigating) and side preference exceeding three standard deviations from the mean.

#### Mounting Behavior

Sexually inexperienced VEH- and DSS-treated male mice were paired with a novel sexually experienced C57BL/6 female mouse in an empty 35 cm x 20 cm x 15 cm (L x W x H) clear plastic box. ANY-maze (Stoelting) was used to record a 15-minute free interaction period. Videos were hand-scored for the frequency and duration of male mounting by a researcher blind to treatment conditions. Mounting behavior was defined as male contact with two paws on the front or back of the female as previously described (Bayless et al., 2023).

#### Open Field Test

The open field was performed in an open-topped grey Plexiglass arena (65 cm x 65 cm x 40 cm; L x W x H). Mice were placed in the centre of the arena and given 20 minutes to freely explore. ANY-maze tracked the total distance travelled and the amount of time spent in the inner and outer zones. The outer zone was defined at 8 cm from the wall of the apparatus.

#### Elevated Plus Maze

The plus-shaped apparatus was elevated 50 cm above the floor and had two open arms and two closed arms, both 30 cm x 8 cm (L x W). The closed arms were surrounded by 15 cm high walls. Mice were placed in the centre of the apparatus and allowed to freely explore for 5 minutes. ANY-maze tracked the total distance travelled and the amount of time spent in the open arms and closed arms.

#### Three Chamber Test

The test apparatus was an open-topped rectangular box (60 cm x 40 cm x 22 cm; L x W x H) divided lengthwise into three identically sized compartments (Yang et al., 2011). The task consisted of three 10-minute phases: habituation, sociability, and social novelty. The sociability and social novelty phases require sex and age matched stimulus mice. The novel mouse was not a cage mate of the familiar mouse in the social novelty phase. Prior to the task, stimulus mice spent 20 minutes/day for 3 days habituating to sitting under inverted pencil cups.

ANY-maze was used to track distance travelled, time in each chamber, and time spent investigating the cup in each phase of the task. When the mouse’s nose was within 1 cm of the pencil cup and pointing directly at it, investigation was scored. Looking above or lateral to the cup and climbing on top of the object were not counted as investigation.

#### Olfaction Habituation-Dishabituation Test

The olfaction test was performed as previously described (Yang and Crawley, 2009). Briefly, mice were habituated to a clean cage without bedding for 30 minutes prior to being repeatedly presented a cotton swab with non-social (vanilla or banana extract diluted 1:1000 in water) and social odours (swab of cage bottom of novel mouse). The order of odour presentation was: water, vanilla extract, banana extract, same sex mice, opposite sex mice. Videos were recorded and hand-scored by researchers blinded to experimental conditions for the time spent investigating each odoured cotton swab, defined as the nose within 1 cm of the swab. Biting and otherwise playing with the cotton swab was not counted as olfaction investigation.

#### Object Location Memory (OLM) Test

The OLM test was performed in an open-topped Plexiglass square box (40 cm x 40 cm x 40 cm; L x W x H) lined with bedding ∼1 cm in depth. Bedding was kept in the same box each day. One wall of the box was lined with duct tape as a visual cue. The OLM took 8 days in total with 6 days of habituation, 1 day of training and 1 day of testing (Vogel-Ciernia and Wood, 2014). Identical cylindrical spice tins filled with cement were used as training objects and were cleaned with diluted saber solution between trials and allowed to fully dry before re-use. The 5-minute testing phase took place 24 hours after training where one object was moved to a novel location and the other remained in the original training location.

ANY-maze tracked the total distance travelled on all 8 days. ANY-maze tracked investigation time and the number of investigations of each object on training and testing days. Investigation time was scored when the mouse’s nose was within 1 cm of the object and pointing directly at the object. Looking over or past the object and climbing on top of the object were not counted as investigation. For the training and testing days, a discrimination index (DI) was calculated where DI is the difference in time spent investigating the moved object and the unmoved object divided by the total time spent investigating both objects x100.

Mice who had a DI over 20 or under −20 during the training phase were determined as having an innate preference and excluded from final analysis. Mice who explored the objects more or less than 3 standard deviations from the mean were also excluded from analysis as we have done previously (Vogel-Ciernia and Wood, 2014).

#### Self-Grooming and Forced Grooming

Mice were individually placed in a new cage without bedding. After a 5 minute habituation period, the cumulative time spent naturally self-grooming was recorded for 5 minutes. The mice were then lightly sprayed with water from a mist bottle and forced self-grooming was scored in the following 5 minutes. All hand scoring of grooming behavior was performed by a researcher blinded to experimental conditions.

#### Forced Swim Test

Mice were placed individually into 1 L glass beakers filled with 900 mL of water at a temperature of 23-25°C. Mice were recorded for 6 minutes and then removed from the water as previously described(Can et al., 2011). The last 4 minutes of footage was hand-scored for time spent immobile, defined as the absence of escape-related movements. The hand-scorer was blinded to experimental conditions.

### Sex steroid quantification

Serum (10 µl) was pipetted into 2 mL polypropylene bead ruptor tubes with 5 ceramic zirconium oxide beads (Hamden et al., 2021, 2019). Calibration curves (0.05 – 1000 pg), blanks, and quality controls were extracted alongside samples via liquid-liquid extraction and analyzed by specific and ultrasensitive LC-MS/MS as before (Hamden et al., 2021). Progesterone, testosterone (T), androstenedione (AE), 5α-dihydrotestosterone (DHT), and 17β-estradiol (E_2_) were measured using multiple reaction monitoring with 2 mass transitions for each analyte. Deuterated internal standards (progesterone-d9, testosterone-d5, and 17β-estradiol-d4) were included in all standards, blanks, controls, and samples to correct for analyte loss and matrix effects. Steroid concentrations were acquired by a Sciex 6500 Qtrap triple quadrupole tandem mass spectrometer. A steroid was considered non-detectable if either the quantifier or qualifier transition was not present. If 50% or more of the values in a group were detectable, then missing values were replaced with the lower limit of quantification ⁄ √2(Handelsman and Ly, 2019) (Handelsman and Ly 2019). DHT and E_2_ were non-detectable in all samples, and AE was non-detectable in all female samples.

### Brain microglia immunofluorescence staining

After PFA fixation and sucrose cryoprotection, brain hemispheres were embedded in Optimal Cutting Temperature (OCT) compound (Scigen, 23-730-625), cryosectioned at 30 μm, and stored in 1X PBS at 4°C. For staining, free-floating sections were permeabilized with 1X PBS + 0.5% Triton X-100 for 5 minutes and then washed 3 x 5 minutes with 1X PBS. Sections were placed in a blocking solution (5% normal donkey serum, and 1% bovine serum albumin dissolved in 1X PBS + 0.03% Triton) for 1 hour at room temperature with gentle shaking and then incubated with anti-IBA1 primary antibody (chicken; Synaptic Systems (234009); 1:1000) for 48 hours. Sections were washed 3 x 5 minutes in 1X PBS + 0.03% Triton and then incubated in a donkey anti-chicken secondary antibody (Alexa Fluor® 488 Jackson ImmunoResearch (703545185); 1:500) and DAPI (Biolegend (422801); 1:1000) for two hours at room temperature on a gentle tilt. Sections were washed 3 x 5 minutes in 1X PBS + 0.03% Triton and 1x 5 minutes in 1X PBS and then mounted onto glass slides and coverslipped using Invitrogen ProLong Glass Antifade Mountant (Invitrogen, P36984).

### Reverse transcription quantitative polymerase chain reaction

At time of euthanasia, the hippocampus tissue was dissected, snap frozen on dry ice and stored at −80°C until RNA extraction. Total RNA was isolated and water blanks using a Monarch® Total RNA Miniprep Kit (New England Biolabs, Whitby, ON) and RNA quantity and quality were confirmed by Nanodrop spectrophotometer. Complementary DNA (cDNA) was reverse transcribed using a LunaScript RT Supermix Kit (New England Biolabs, Ipswich, MA). The amount of RNA used for reverse transcription was normalized (227 – 400ng) and experimenters were blinded to treatment group and sex during RNA isolation and cDNA synthesis. Transcripts (*Tlr4, Nod1, Nod2, Il1β, Tnfα, Il6*) were quantified via SYBR Green assays (Integrated DNA Technologies, Inc., Coralville, IA) (Table 1). Relative expression levels of the reference gene and genes of interest were assessed using Luna® Universal qPCR Master Mix (New England Biolabs, Whitby, ON). Samples were run in duplicate. A reference gene (*Hprt*) was measured in each sample. Negative controls (water blanks, no reverse transcriptase controls) confirmed the specificity of assays. Melt curve analyses verified that the expected targets were amplified. One sample was excluded from the Male DSS *Nod1* analysis due to lack of specificity in the melt curve.

**Table 1.**
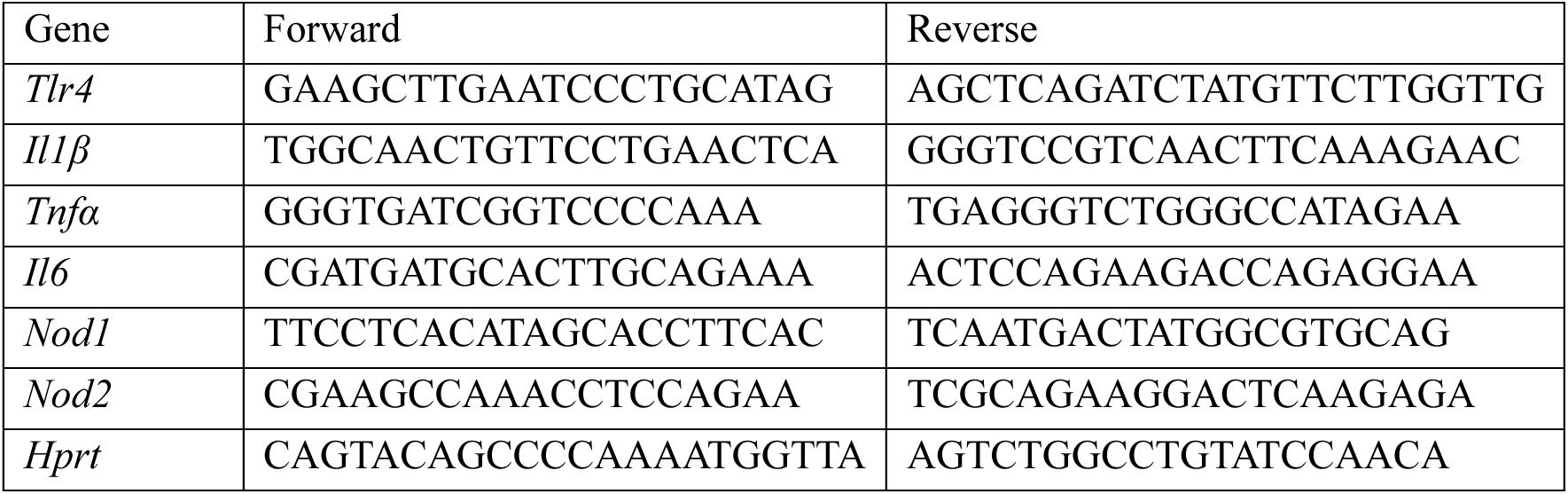
RT-qPCR primers.

The cycle threshold (Ct) of the reference gene *Hprt* for each treatment group and sex were analyzed by two-tailed t-test to determine whether expression of reference gene differed between VEH and LPS groups. There was no significant difference between VEH and DSS groups for the females nor males (p = .76 and .31, respectively). The mean of the reference gene was used to calculate the ΔCq and ΔCq values were normalized to the VEH group within each sex. Statistics were run and data were graphed using the fold change for each transcript within each sex. Data were analyzed for an effect of treatment by unpaired t-tests.

### Microglial morphology analysis

Slides were imaged using the Zeiss Axioscan7 slide scanner microscope at 20X magnification. Maximum intensity extended depth of focus (EDF) images were acquired from Z-stacks containing 16 slices at a step size of 1 µm. EDF images were created using maximum projection settings and converted to tiffs using ZEN (Zeiss). Maximum projection EDF images compile the pixels of highest intensity at any given z-stack position and construct a new 2D image retaining the 3D information. The hippocampus and hypothalamus anatomical regions were traced using the FASTMAP FIJI plugin with reference to the Allen Brain Atlas (Terstege et al., 2022). Within these segmented regions, microglial morphology was analyzed using the pipeline described in Kim et al., 2024 using MicrogliaMorphology (ImageJ tool) and MicrogliaMorphologyR (R package) (Kim et al., 2024a) which uses Image-J plugins Skeletonize (2D/3D) and AnalyzeSkeleton (2D/3D) (Arganda-Carreras et al., 2010) to collect 27 metrics of morphology per microglial cell (Supplemental Figure 4A). We conducted dimensionality reduction using principal component analysis followed by k-means clustering on the first three principal components (Supplemental Figure 4C) to define four morphological classes within our dataset (hypertrophic, rod-like, ramified, ameboid). We identified the optimal clustering parameter using exploratory data analysis methods including the within sum of squares and silhouette methods (Supplemental Figure 4B). We assigned cluster identities by examining the relationship between each cluster and the 27 morphological features (Supplemental Figure 4D) and by using the ColorByCluster feature in MicrogliaMorphology to visually examine the different cluster morphologies (Figure 7A).

We then calculated the percentage of cells in each morphology cluster for each treatment and sex using the ‘clusterpercentage’ function within MicrogliaMorphologyR. To assess how cluster membership changes with treatment for each sex, we fit a generalized linear mixed model using a beta distribution to model the percentage of cluster membership as a factor of Cluster identity and DSS treatment with MouseID as a repeated measure ("percentage ∼ Cluster*treatment + (1|MouseID)") using the ‘stats_cluster.animal’ function from MicrogliaMorphologyR, which is wrapped around the glmmTMB R package (Brooks et al., 2017). We ran a 2-way Analysis of Deviance (Type II Wald chisquare test) test on the model to assess the contribution of Cluster and treatment interactions. Tests between treatment conditions were corrected for multiple comparisons across Clusters (∼treatment|Cluster) using the Sidak method and q-values < 0.05 were considered statistically significant.

### Short and medium chain fatty acid quantification

Serum samples were stored in −80 freezer prior to lipid extraction. A total of 50 µL of serum was mixed with 450 µL ice-cold methanol, vortexed for 10 seconds, and incubated at −20 °C for 4 hours for protein precipitation. Samples were then centrifuged at 14,000 rpm at 4 °C for 15 minutes, and the supernatant were collected and dried down via SpeedVac at 4 °C. The dried extract was reconstituted in 100 µL of water and acetonitrile solution (1:1, v:v) for LC-MS analysis. Quality control (QC) samples were prepared by pooling 20 μL aliquot from each sample to monitor the instrument performance and optimize the injection volume. A method blank sample was prepared using the identical analytical workflow but without serum. An UHR-

QqTOF (Ultra-High Resolution Qq-Time-Of-Flight) Impact II (Bruker Daltonics, Bremen, Germany) mass spectrometry interfaced with an Agilent 1290 Infinity II Ultrahigh-Performance Liquid Chromatography (UHPLC) system (Agilent Technologies, Santa Clara, CA, USA) was used for analysis. LC separation was achieved using a Waters reversed phase (RP) UPLC Acquity BEH C18 Column (1.7 µm, 1.0 mm ×100 mm, 130 Å) (Milford, MA, USA) maintained at 25 °C. The mobile phase A was 5% ACN in H_2_O with 10 mM ammonium acetate (pH = 9.8, adjusted ammonium hydroxide) and the mobile phase B was 95% ACN in H_2_O. The LC gradient was set as follow: 0 min, 5% B; 8 min, 40% B; 14 min, 70% B; 20 min, 95% B; 23 min, 95% B; 24 min, 5% B; 33 min, 5% B. The flow rate was 0.1 mL/min. Negative ion mode was applied with the following MS settings: dry gas temperature, 220 °C; dry gas flow, 7 L/min; nebulizer gas pressure, 1.6 bar; capillary voltage, 3000 V. External calibration was applied using sodium formate to ensure the *m/z* accuracy before sample analysis.

SCFA and MCFA measurements from cecal contents were obtained as follows. In brief, 20-40 mg of cecal contents were mixed with 0.4 – 0.8 mL of 25% phosphoric acid in water (LabChem). Sample mixtures were thoroughly homogenized via vortexing, then centrifuged at 15,000 x g for 10 minutes at 4°C. Following centrifugation, the sample supernatant was transferred to a new tube, then centrifuged again at 15,000 x g for 10 minutes at 4°C. The resulting supernatant was then frozen at −20°C before being sent to the University of Alberta Agricultural, Food & Nutritional Science (AFNS) Chromatography Facility for volatile fatty acid (VFA) analysis using gas chromatography/mass spectrometry (GC/MS) as previously described (Ng et al., 2023).

There, the supernatant was filtered through 0.45 µm filters and mixed in a 4:1 ratio with internal standard solution (24.5 mmol/L isocaproic acid). The samples were then injected into a Stabilwax-DA column (length: 30 m, inner diameter: 0.53 mm, film thickness: 0.5 µm, Restek Corporation) on a Bruker Scion 456 GC with a model 8400 autosampler (Bruker Ltd) using a helium carrier gas. The samples were run using a 10 mL/min constant flow with a 5:1 split ratio with a column temperature gradient as follows: 80°C initial temperature, immediately increased to 180°C at 20°C/min, then held for 3 min at 180°C. The injector and detector temperature were at 250°C. The peaks were analyzed using CompassCDS software. The concentration of SCFAs/MCFAs per sample was then normalized to mass of the cecal contents for statistical testing.

### Bacterial culturing and preparation for SCFA quantification

Selected bacterial strains (Table 2) from public culture collections and lab strain libraries were cultured anaerobically in a vinyl anaerobic chamber maintained with an atmosphere of 5% CO_2_, 5% H_2_, and 90% N_2_ for 1-5 days (Coy Lab Products, Grass Lake, MI, USA) on pre-reduced Anaerobic Akkermansia Media plates (AAM:18.5 g/L Brain Heart Infusion, 5.0 g/L Yeast Extract, 8.5 g/L Tryptone, 1.5 g/L Soytone, 2.5 g/L NaCl, 3.75 g/L K_2_HPO_4_, 1.75 g/L *D*-glucose, 0.4 g/L Na_2_CO_3_, 1.0 mg/L Hemin, 0.5 g/L L-Cysteine HCl, 0.5 mg/L Vitamin K1, and 3% v/v heat-inactivated fetal calf serum supplemented with 15 g/L Agar) until visible colonies were observed. Subsequently, strains were inoculated in AAM for 3 days (or 1 day for fast-growing strains) before being centrifuged at 6000 xg for 5 minutes to remove bacteria. 1 mL of bacterial supernatants was then combined with 200 µl of 25% v/v phosphoric acid before being frozen and quantified at the AFNS Chromatography Facility as described above.

**Table 2.**
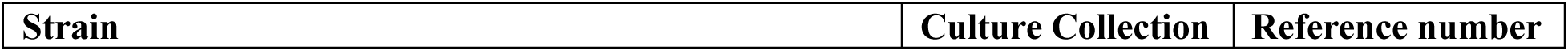
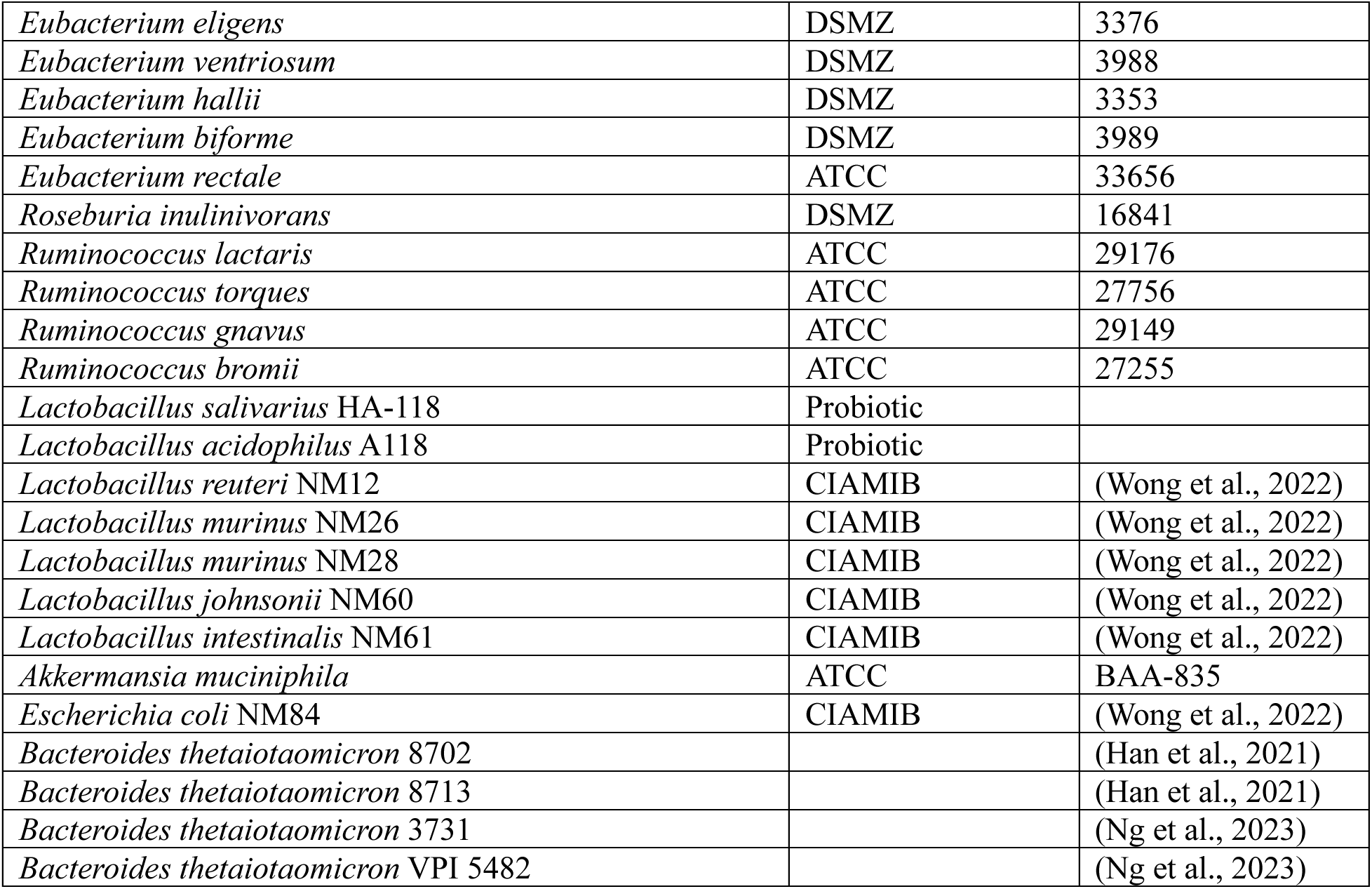
Cultured bacterial strains used for *in vitro* SCFA measurements.

### Bacterial metagenomics shotgun sequencing and analysis

DNA from cecal contents collected from mice 3 days after cessation of the third round of DSS was extracted and quantified as described above in “*Microbiome sequencing and analysis”*. Shotgun metagenomics library preparation was done using the Illumina DNA Prep library preparation kit in accordance with manufacturer instructions, and sequencing was then performed on the Illumina NextSeq^TM^ platform with 2×150 bp paired-end read chemistry. Library preparation and sequencing were performed at the UBC Sequencing + Bioinformatics Consortium.

For shotgun sequencing analysis, the quality of raw reads was evaluated using FASTQC v0 12.1 (Andrews, 2010) and MultiQC v 1.19 (Ewels et al., 2016). Data was then run through SqueezeMeta v 1.6.5 pipeline in coassembly mode (Tamames and Puente-Sánchez, 2019). The coassembly method pools reads from all samples to create a single shared metagenome assembly. Briefly, sequences were trimmed and quality-filtered using Trimmomatic (Bolger et al., 2014); metagenome assembly was performed using Megahit (Li et al., 2015); ORF and rRNA prediction was done using Prodigal (Hyatt et al., 2010) and barrnap (Seemann, 2024), respectively before classification was performed using RDP classifier (Wang et al., 2007); Diamond software (Buchfink et al., 2015) was used to assign taxonomy against the Genbank nr database, COGs/NOGs against the eggNOG database (Huerta-Cepas et al., 2016) and KEGG ID against the KEGG database (Kanehisa and Goto, 2000); gene coverage and abundance estimation were performed by mapping individual reads to the reference coassembly contig using Bowtie2 (Langmead and Salzberg, 2012). Finally, SqueezeMeta outputs were analyzed using the SQMTools package (Puente-Sánchez et al., 2020). Normalization of gene abundance was performed using trimmed mean of M-values (TMM) with the edgeR package (Robinson et al., 2009) and plotted using ComplexHeatmap (Gu et al., 2016). The TMM method of normalization can robustly account for differences in sequencing depth between samples by adjusting raw counts using a unique scaling factor for each sample. The scaling factor is derived by comparing gene abundances against a reference sample (which is automatically determined to be the sample that has the 75^th^ percentile of log-counts closest to the median across all samples), then using a weighted trimmed mean over the log-transformed fold-change differences between the sample and the reference.

### β-glucuronidase assay

Beta-glucuronidase activity was measured using the QuantiChrom^TM^ β-Glucuronidase Assay Kit (BioAssay Systems, San Francisco, CA, USA). Cecal contents collected from mice acutely following DSS (P71) and after ∼40 days of recovery (P110) were processed as follows: 10-50 mg of cecal contents were aliquoted and suspended in 600 µl of sterile dH_2_O. Samples were then homogenized at 30 Hz for 2 minutes using the TissueLyser II (QIAGEN) before being centrifuged at 1000 xg for 1 minute at 4°C to remove cecal debris without pelleting bacteria. The supernatant was then centrifuged at 13000 xg for 5 minutes at 4°C. The pelleted bacteria were then resuspended in 600 µl of chilled sterile dH_2_O and vortexed for 10 seconds. Bacterial lysis was achieved through 3 rounds of sonication at 50 watts for 20 seconds each using an XL2020 sonicator equipped with a Microtip^TM^ (Misonix, Farmingdale, NY, USA). Samples were diluted 1:2-1:10 as needed before performing the β-glucuronidase activity assay according to manufacturer instructions. Enzyme activity was normalized to total protein concentration in each sample, measured using the Pierce BCA Protein Assay Kit (ThermoFisher).

### MRI acquisition and data processing

*Ex vivo* imaging and analysis, including standard data preprocessing and region of interest (ROI) analyses, were performed as previously described (Yi et al., 2019). Briefly, brains were extracted from the cranial vault between P95 and P130 and post fixed in 4% PFA (n = 54;13 vehicle per sex, 14 DSS per sex). Brains were placed in a custom-built holder immersed in Fluorinert (FC-3283, 3M, St. Paul, MN, USA) and imaged with a 4.7-T Agilent MRI system with a 3.5 cm diameter quadrature volume RF coil. Multi-slice, diffusion-weighted, spin echo images were used to acquire 10 nondiffusion weighted images (b=0 s•mm^−2^) and 75 diffusion-weighted images (25: b=800 s•mm^−2^, 50: b=2,000 s•mm^−2^), using non-colinear diffusion-weighting directions. Diffusion imaging was performed with a TE/TR=24.17/2000-ms, FOV=30×30 mm^2^, and a matrix=192×192 reconstructed to 256×256 for an isotropic voxel size of 0.25-mm over two signal averages. Raw data files were converted to NIfTI format and FSL was used to correct for eddy current artifacts (Smith et al., 2004). A DTI-based mouse brain atlas (Jiang and Johnson, 2011) was used as a study space template and to define regions of interest (ROI) including the left and right hippocampus, frontal association cortex, striatum, amygdala, thalamus, and hypothalamus. Multi-shell diffusion data were fit with the Microstructure Diffusion Toolbox (MDT) (Harms et al., 2017) to the neurite orientation dispersion and density imaging (NODDI) model (Zhang et al., 2012). An additional compartment of isotropic restriction was included to account for potential fixative effects as recommended (Alexander et al., 2010). The mean values for neurite density index (NDI) and orientation dispersion index (ODI), which represent indices of intracellular and extracellular diffusion respectively, were calculated in each ROI using Pyradiomics (van Griethuysen et al., 2017) and compared among experimental groups.

### Statistical analyses

Statistics and graphs were created with GraphPad Prism 10.1.1 (GraphPad Software), Python 3.8 and RStudio software (RStudio Team, 2022). A repeated measures ANOVA was used to assess body weight changes during treatment. All organ comparisons, SCFA measurements, gut microbial β-glucuronidase (GUS) activity measurements, and almost all behavioral tests were assessed using a two-way ANOVA to assess main effects of treatment and sex and Šidák corrected post-hoc comparisons were performed to determine statistical significance within sex if main effects or interactions were observed. The urine preference test was assessed by comparing each group to chance (50%) using a single sample t-test. To directly compare preferences between treatments, an unpaired two-tailed t-test was conducted. Steroid levels were analyzed by Mann-Whitney U. Statistical significance of alpha diversity metrics (Observed features) was assessed using a two-way ANOVA to assess main effects of treatment and sex. Beta diversity metrics were evaluated using PERMANOVA analysis. FDR-corrected p-values obtained from DAA with metagenomeseq were reported.

For the MRI analysis, a 2 × 2 analysis of variance (ANOVA) was used to determine the effect of treatment, biological sex, and their interactions on the ROI values. The effects of ROI laterality and imaging age effects were regressed from the data before the ANOVA was performed as these effects were not pertinent to the tested hypotheses. Benjamini-Hochberg false discovery rate (FDR) procedure was used to control for multiple hypothesis testing across ROIs within each imaging signal. Statistical significance of an ANOVA is defined as an FDR adjusted p-value <.05. Post-hoc pairwise difference testing was performed using Fisher’s LSD test Partial η^2^ explains how much of the unexplained variance in the model is described by the addition of that variable into the model. The plotted imaging metric values represent the metric values after the laterality and batch effects have been removed.

## Results

### An early-life DSS model of pediatric IBD

To establish a model of juvenile-onset IBD that mimics the remitting and recurrent cycles of gut inflammation often observed in juvenile patients with IBD, we administered DSS in drinking water repeatedly across early postnatal development starting on postnatal day 21 (P21) (Figure 1A). Colon length is indicative of inflammatory state, with colon shortening being induced by effective DSS treatment (Chassaing et al., 2014). Three days after the final treatment, colon length was significantly decreased by DSS in both sexes (main effect of treatment (F (1, 21) = 39.30, *p*.0001) (Figure 1B). Both female (*p*=.009) and male (*p*=.0002) mice treated with DSS had colon lengths that were significantly shorter than untreated mice, indicating successful induction of intestinal inflammation in our model. Furthermore, body weights were recorded throughout the treatment period (Chassaing et al., 2014) (Figures 1C, D). Female mice showed mild weight loss in response to DSS, with only the second round producing significant decreases from VEH littermates (treatment x day interaction F(50,714) = 6.693, *p*<.0001) (Figure 1C). Conversely, male DSS-treated mice showed more dramatic weight loss across the three treatment periods (treatment x day interaction F(50,686) = 12.372, *p*<.0001) (Figure 1D). To further investigate inflammation caused by repeated administration of DSS, we examined the histological architecture of colonic tissues using hematoxylin and eosin staining. Following chronic DSS treatment, the gut epithelial layer was disturbed, showing increased crypt length (crypt abscess), depletion of mucus-producing goblet cells, increased incidence of inflammatory cell infiltration, and ruffling of the normally smooth epithelium (Figures 1E, F). Furthermore, UEA-1 lectin staining targeting the intestinal mucus layer revealed that DSS-treated juvenile mice experienced profound alterations in mucus structure, with loss of the normal striated bilayer organization (Ng and Tropini, 2021) (Figure 1F). Finally, to examine changes to gut microbiota composition caused by repeated cycles of early life inflammation, we performed 16S rRNA sequencing targeting the bacterial V4-V5 variable regions. Consistent with previous studies in adult mice, DSS treatment resulted in significant losses to microbiota alpha diversity, with the number of observed features, a proxy for the number of bacterial species present (Kleine Bardenhorst et al., 2022), being significantly decreased in DSS-treated animals compared to vehicle (significant effect of treatment F(1,12) = 5.3, *p* = .0387) (Figure 1G) (Munyaka et al., 2016). Similarly, between-group microbiota compositional differences were revealed to be significantly different when compared using calculated Bray-Curtis distance matrices (treatment x sex interaction F(1,12) = 3.2158, *p* < .046) (Figure 1H). Altogether, these results suggest that our DSS colitis model of juvenile IBD induces profound physiological and microbiological changes to the gut.

### Early-life DSS affects self-grooming behavior in male mice

Previous work in early life gut-inflammatory models, including those for IBD, identified altered mouse behavior (Salvo et al., 2020). To fully profile behavioral phenotypes in our pediatric IBD model, we performed a battery of behavioral testing beginning 17 days after DSS treatment to examine anxiety-like (elevated plus maze, light-dark box, open field test), memory (spatial object location), sociability and social memory (three chamber task), repetitive behavior (self-grooming), and depression-like behaviors (forced swim). Unlike previous reports, we did not observe any significant effects of treatment for either sex in most of these tests (Figure 2 and Supplemental Table 1), with the single exception of decreased self-grooming (main effect of treatment (F (1, 24) = 6.514, *p*=.018)) in the DSS-treated males (*p*=.031). There were no significant main effects of treatment or interactions with treatment on distance travelled or total exploration on any of the tasks (Supplemental Figure 1 and Supplemental Table 1). This suggests that that the majority of behaviors examined are not impacted by previous repeated bouts of postnatal gut inflammation in early-life, potentially due to the normal development of most neuronal circuits prior to treatment beginning at P21.

**Figure 2.**
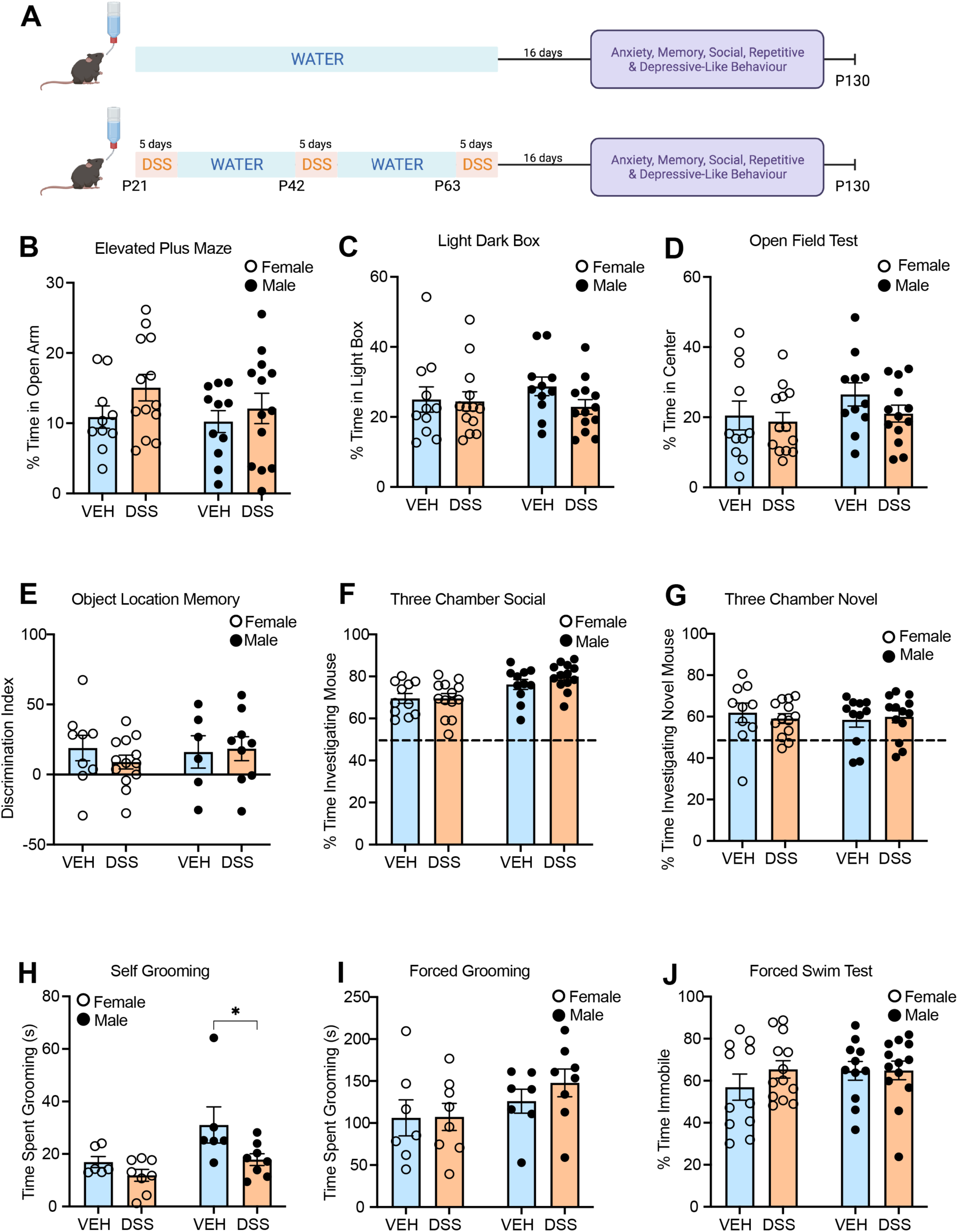
Early-life DSS treatment has minimal impacts on adult anxiety, social, memory, and depression-like behaviors. *A*. Experimental design. After a 16-day recovery period, a behavioral battery compared VEH and DSS treated mice on *B.* the elevated plus maze (n=10-13/group) *C.* the light dark box (n=11-13/group) *D.* the open field test (n=11-13/group) *E*. the object location memory test (24hr) (n=6-13/group) *F*. the three-chamber social task (n=11-13/group) *G.* the three-chamber novel social task (n=10-13/group) *H.* self-grooming (n=6-8/group) *I*. forced grooming (n=7-8/group) and *J.* the forced swim test (n=11-13/group). All graph bars and error bars represent the mean ± SEM. \**p*<.05

### Early-life DSS disrupts male preference for female odour

To test the impact of repeated bouts of inflammation on sexual behaviors in our model, ∼1 month following DSS, mice were given a preference test for same- or opposite-sex urine (Figure 3A). Female VEH- and DSS-treated mice both failed to show a significant preference for either same-sex urine (VEH one sample t-test vs 50% *t*(6) =0.383, *p*=.714 or DSS one sample t-test vs 50% *t*(6) =0.406, *p*=.70) or opposite-sex urine (VEH one sample t-test vs 50% *t*(6) =0.350, *p*=.74 or one sample t-test vs 50% DSS *t*(6) =0.256, *p*=.81) compared to water (Figure 3B). In contrast, male VEH-treated mice showed a significant preference for same-sex urine compared to water (one sample t-test vs 50% *t*(7) = 2.468, *p*=.04) and a trend for a preference for opposite-sex urine *t*(7) = 1.972, *p*=.086. However, male DSS-treated mice failed to show a preference for same-sex urine (*t*(7) = 1.153, *p*=.29) as well as opposite-sex urine over water (*t*(7) = 0.99, *p*=.36) (Figure 3C). When comparing preferences directly, there was no significant difference in VEH-treated and DSS-treated males in their preference for male urine (*t*(14) = 1.45, *p*=.17). However, VEH-treated males showed a significantly higher preference for female urine than DSS-treated males (*t*(14) = 2.16, *p*=.048) (Figure 3C). Together, these data demonstrate that DSS treatment produced a significant deficit in opposite-sex scent-seeking behaviors, a key aspect of mate-seeking behavior in male rodents (Malkesman et al., 2010).

**Figure 3.**
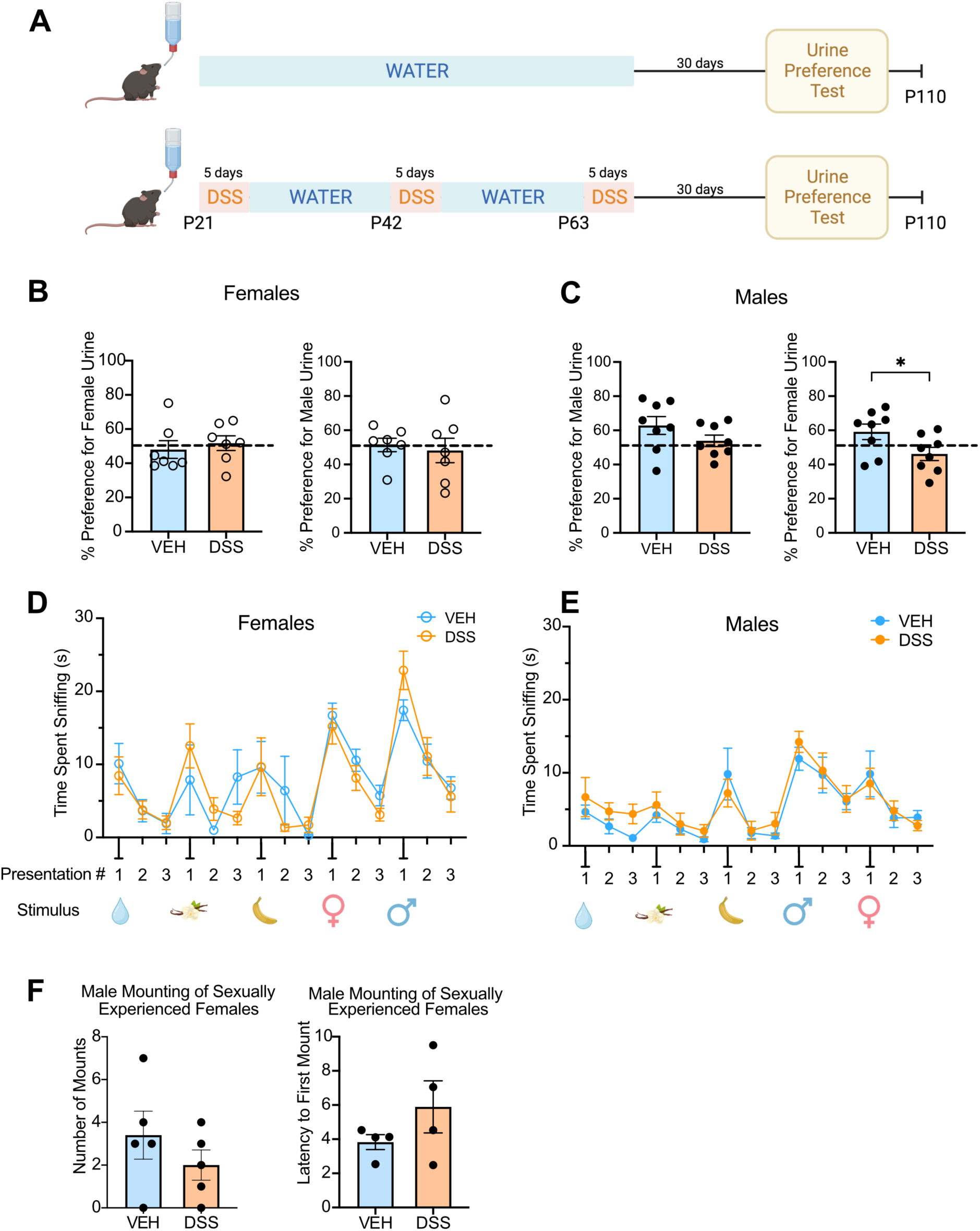
Early-life DSS treatment induces male sex-specific impairments in mate-seeking behavior. *A*. Experimental design. After a 30-day recovery period, mate-seeking behavior was assessed by the Urine Preference Test. *B*. Female mice showed no preference for female or male urine over water and VEH and DSS female mice performed similarly. n=7/group. Each graph bar and error bar represent the mean ± SEM. Dotted line is 50% preference. *C*. Male VEH mice significantly preferred both male and female urine over water (above 50% dotted line). DSS treated males showed no preference for urine over water. n=8/group. *D, E.* Olfactory habituation-dishabituation test for females (*D*) and males (*E*) revealed no differences between DSS- and VEH-treated mice and normal exploration, habituation and then dishabituation patterns for each odour n=6-7/group. *F.* DSS and VEH male mice showed similar number of mounts and latency to first mount a sexually experienced female. n=4-5/group. \**p*<0.05.

To verify that deficits in the urine preference test in DSS-treated males were not due to impairments in olfaction, we performed the olfaction habituation-dishabituation task (Yang and Crawley, 2009)We tested two extract odours (vanilla and banana) and two social odours (one same sex and one opposite sex novel mouse odours). Both treatment groups and sexes showed the expected patterns of active exploration of each odour followed by habituation across repeated trials and then active exploration of the next novel odour (dishabituation) (Figure 3D and E and supplemental Table 1), indicating that DSS treatment did not negatively impact odour processing or recognition. Together, these findings demonstrate a long-term shift in male-specific preferences for female odours following early-life gut inflammation.

To further explore impacts on consummatory mating behaviors, we compared sexually naïve mounting behavior of the VEH- and DSS-treated male mice when paired with a sexually experienced untreated female wildtype control mouse. Over a 15-minute mating session, DSS-treated male mice showed no significant differences in copulatory behaviors compared to VEH males including frequency of mounting (*t*(8) = 1.06, *p*=.32) and latency to first mount (*t*(6) = 1.30, *p*=.42) (Figure 3F). Together, this suggests that early-life DSS treatment has specific deficits on mate-seeking through olfactory cues and not on consummatory sexual behavior.

### Early-life intestinal inflammation disrupts androgens in males

Given the observed differences in urine odor preferences, we investigated reproductive organ development in DSS- and VEH-treated mice. Male and female reproductive organs were collected following behavior testing and weighed (P110). In males, there was a significant decrease in the weight of the androgen-sensitive seminal vesicles in the DSS-treated animals compared to VEH (*t*(23) = 2.44, *p*=.03) (Figure 4A). However, there was no significant difference in testes weight between treatments (*t*(23) = 1.05, *p*=.30) (Figure 4B). In females, we observed no difference in uterus weight (*t*(23) = 0.42, *p*=.68) (Figure 4C). To further investigate the impairment in male seminal vesicle development, we examined changes in circulating steroid hormones in VEH- and DSS-treated male mice both acutely (P71, 3 days after final DSS treatment) (Figure 4D, E) and chronically (after behavioral testing, P110) (Figure 4F, G). At the acute timepoint (P71), the DSS-treated males showed a significant decrease in serum testosterone (U = 0, *p =*.03) and androstenedione (U = 0, *p =*.03) levels. This decrease was maintained through behavioral testing for androstenedione (U = 27, *p* =.01) but not for testosterone (U = 46, *p* =.22). There were no significant differences in serum progesterone in males at either timepoint and no impacts in females for androstenedione, testosterone, or progesterone (Supplemental Figure 2). All together, these data indicate that early-life DSS treatment reduces male reproductive organs, potentially through reduced levels of circulating androgens.

**Figure 4.**
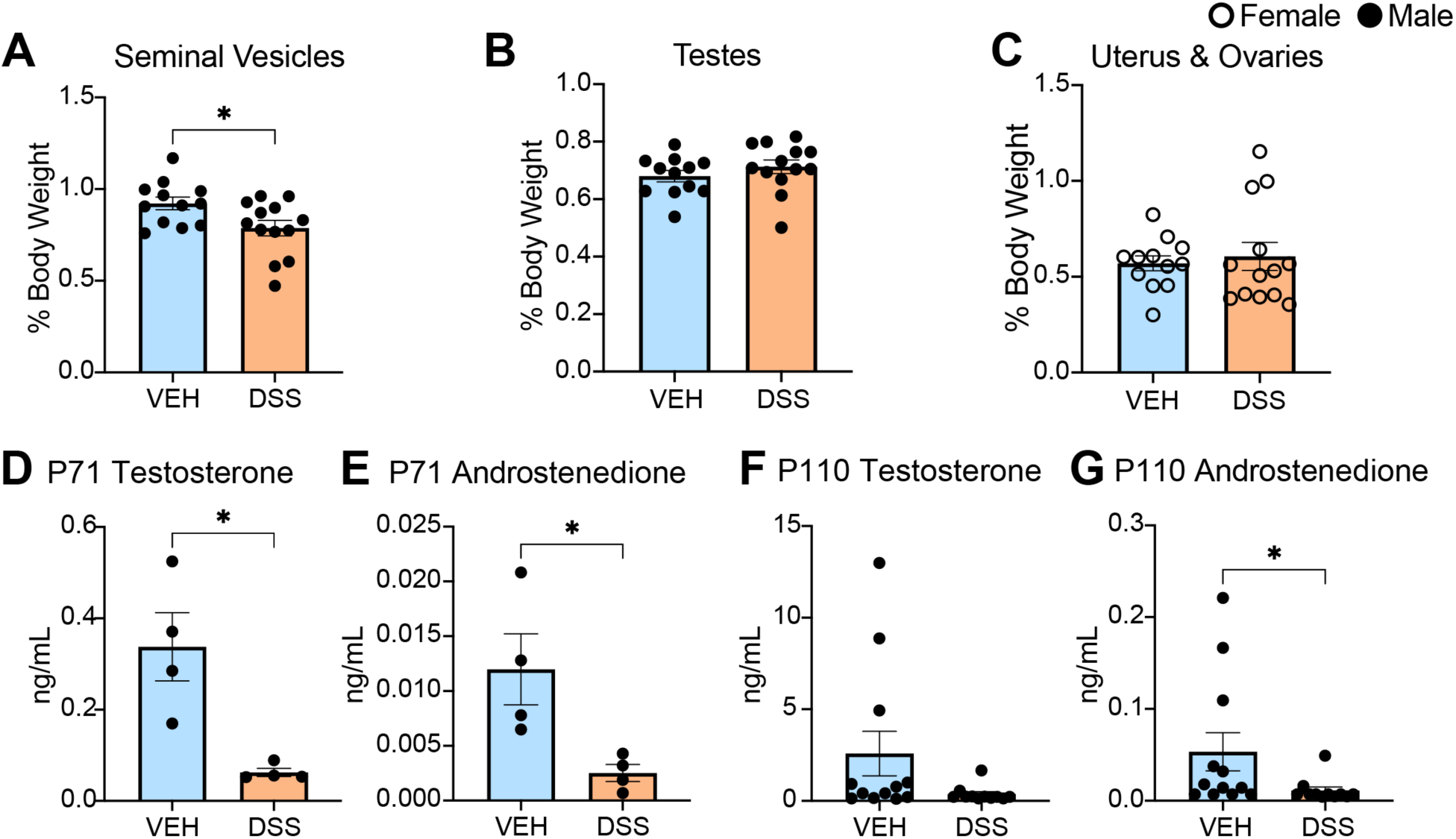
Early-life DSS treatment induces male sex-specific impairments in reproductive physiology. *A*. Seminal vesicles were significantly lower in weight in DSS-treated male mice than VEH-treated mice at P110. n=12-13/group. *B*. There were no significant differences in testes weight between DSS- and VEH-treated males at P110. n=12-13/group. *C*. No significant differences in uterus and ovaries weights between DSS- and VEH-treated females were observed at P110. n=12-13/group. *D-E*. 3 days after the last DSS treatment (P71), male serum testosterone (*D)* and androstenedione (*E*) levels were significantly lower in DSS-than in VEH-treated mice. n=4/group. *F-G*. Following the last behavioral experiment (P110), male serum testosterone (*D)* levels were not significantly different between VEH and DSS mice, but androstenedione (*E*) levels were significantly lower in DSS-than in VEH-treated mice. n=12-13/group. Each graph bar and error bar represent the mean ± SEM. \**p*<0.05.

### Early-life intestinal inflammation alters SCFA/MCFA production and gut microbiome composition to favor decreased abundance of SCFA/MCFA-producing taxa

Since we observed changes in microbiome composition immediately following DSS treatment, we then examined circulating short and medium-chain fatty acids (SCFAs and MCFAs respectively) (Figure 5A), which are produced by the gut microbiota. Importantly, SCFAs are known to be disrupted in patients with IBD and can alter sex steroids (Acharya et al., 2024) and brain function (Cryan and Dinan, 2012; Erny et al., 2015; Sullivan and Ciernia, 2022). Acutely following the last round of DSS (P71), DSS treatment significantly reduced female (*p* = .014) and male (*p* = .037) serum levels of valeric acid (see Supplemental Table 1 for full statistics). In females, DSS also decreased serum levels of capric acid (*p* = .022) (Figure 5A). Interestingly, we found no significant differences in levels of acetic, propionic, and butyric acid, which are the most abundant SCFAs and play important roles in maintaining intestinal homeostasis (Venegas et al., 2019). We also measured SCFA/MCFA levels in cecal contents, where butyric acid was the only SCFA that was significantly different between VEH- and DSS-treated males (*p* = .0498) (Figure 5B, see Supplemental Table 1 for full statistics). Critically, butyric acid is an important anti-inflammatory agent that is known to be reduced in patients with IBD (Hodgkinson et al., 2023).

**Figure 5.**
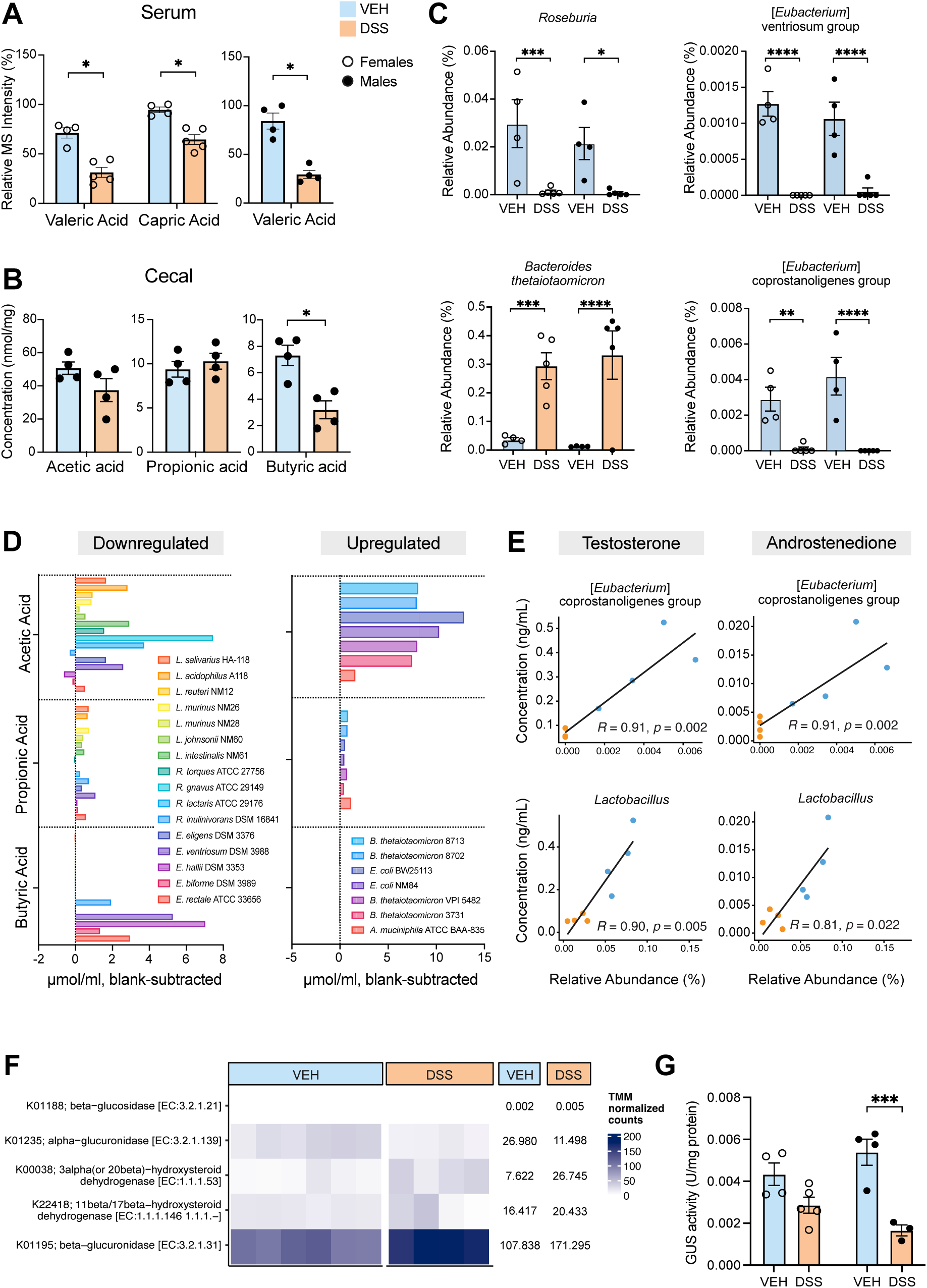
Early-life DSS treatment causes decreases in short- and medium-chain fatty acid levels and relative abundance of associated microbiota. *A*. Serum levels of valeric and caproic acid were significantly lower in DSS-treated females compared to VEH at P71. Valeric acid was also significantly lower in DSS-treated male serum. n=4-5/group. See Supplement for full set of serum statistics. *B.* Cecal levels SCFAs at P71, represented in nmol/mg of contents. Concentration of butyric acid was significantly different in DSS-treated males compared to VEH at P71. n=4-5/group. See Supplement for full set of cecal SCFA statistics. *C.* Relative abundance of select SCFA-producing bacteria. n=4-5/group. *D. In vitro* SCFA production from bacterial genera represented in the experimental microbiota. *E.* Correlation of select bacteria with serum testosterone and androstenedione concentrations in males. n=4/group. *F.* Metagenomic abundance of steroid-modifying and steroid-degrading enzymes normalized by TMM, trimmed mean of M-values. Numbers represent mean TMM-normalized counts per treatment for each enzyme. n=4-6/treatment. *G.* GUS activity in cecal contents was significantly lower in DSS-treated males. Activity was measured from cecal contents and normalized to total protein. n=3-5/group. Error bars in bar plots represent SEM. \**p*<0.05, \*\*\**p*<.001, \*\*\*\**p*<.0001.

We then investigated whether SCFA-producing bacteria were differentially present in VEH- and DSS-treated mice acutely following the last round of DSS at P71 (Figure 5C, Supplemental Figure 3A). Differential abundance analysis revealed that DSS-treated female mice possessed decreased relative abundance of many bacteria genera belonging to the phylum Bacillota (Supplemental Figure 3B, See Supplemental Statistics Table for full list), which contains many SCFA-producing bacterial species (Fusco et al., 2023; Mandal et al., 2015). Indeed, the relative abundance of Bacillota was moderately correlated with cecal acetic (*R* = 0.54, *p* = .005) and butyric acid (*R* = 0.76, *p* < .001) levels (Supplemental Figure 3C). Specifically, DSS-treated mice possessed decreased relative abundance of the genus *Roseburia* (*p_females_* = .0039, *p_males_* = .0201) along with several *Eubacterium* groups ([*Eubacterium*] ventriosum group, *p_females_* < .0001, *p_males_* < .0001; [*Eubacterium*] xylanophilum group, *p_females_* < .0001, *p_males_* = .0062; [*Eubacterium*] coprostanoligenes group, *p_females_* = .0059, *p_males_* < .0001) – bacterial taxa that are known to produce SCFAs (Figure 5C, D, Supplemental Figure 3B) (Akhtar et al., 2021; Fusco et al., 2023; Mirzaei et al., 2021; Ziętek et al., 2021). Differential abundance analysis also revealed that the species *Bacteroides thetaiotaomicron* (*p_females_* = .0001, *p_males_* < .0001) and *Akkermansia muciniphila* (*p_females_* < .0001, *p_males_* = ns) were enriched in DSS-treated mice, bacteria not known to produce butyric acid (Effendi et al., 2022; Kim et al., 2021). To investigate these correlations, we cultured a variety of strains belonging the above identified taxa, then measured production of SCFAs *in vitro* (Figure 5D). This analysis revealed that *Eubacterium* sp. and *Roseburia sp.* are both capable of producing butyric acid and are depleted following DSS treatment, potentially contributing to the decreased concentration of butyric acid in cecal contents. Interestingly, strains that are both more (*Bacteroides* sp. and *Akkermansia* sp.) and less (*Ruminoccocus* sp. and *Lactobacillus* sp.) abundant after DSS treatment were capable of producing acetic acid, which may contribute to the lack of significant difference in cecal acetic acid concentration following DSS.

Due to the differences observed in mating behaviour and circulating sex hormone levels following DSS treatment, we also examined the potential role of the DSS-altered microbiome in driving these phenotypes. Of the differentially abundant taxa observed, we noted that many have been shown to express β-glucuronidases (GUS) and β-glucosidases, enzymes that convert conjugated hormones to their active form (Ervin et al., 2019; Patel et al., 2023). Strikingly, the relative abundance of several GUS-producing taxa was correlated with serum testosterone and androstenedioine concentrations (Figure 5E). For instance, the relative abundance of [*Eubacterium*] *coprostanoligenes* group was positively correlated with androgen levels (androstenedione, *R* = 0.91, *p* = .002; testosterone, *R* = 0.91, *p* = .002). Similarly, the relative abundance of *Lactobacillus* was positively correlated with serum androstenedione (*R* = 0.81, *p* = .022) and testosterone (*R* = 0.90, *p* = .005). In contrast, the relative abundance of *Bacteroides thetaiotaomicron*, which is known to express a sulfotransferase that can deactivate hormones (Cotton et al., 2023), was negatively correlated with testosterone concentrations (*R* = −0.74, *p* = .046) (Supplemental Figure 3D).

Following this observation, we performed shotgun metagenomic sequencing analysis to examine the genomic potential of the microbiota to express hormone-modifying and degrading enzymes (Figure 5F). Supporting the genetic potential of the microbiota in our mice to modify active levels of hormones, we found the presence of genes mapping to various hydroxysteroid dehydrogenases (K00038 and K22418) as well as β-glucuronidase (K01195). Importantly, genomic potential is not a predictor of expression levels or enzyme activity due to various pre- and post-translational regulatory mechanisms. Therefore, we aimed to measure GUS activity directly from cecal contents using a fluorometric molecule conjugated to a glucuronide. Interestingly, GUS activity was significantly decreased in males (*p* = .0006) following DSS treatment, with activity in females trending lower as well (*p* = .0856) (Figure 5G). Taken together, these findings indicate that early life inflammation alters microbiome composition, leading to differences in SCFA producers and GUS activity which correlates with SCFA and androgen levels.

### Early life intestinal inflammation causes long-lasting physiological differences and changes to gut microbiota composition

Having observed changes in microbially-produced and modified metabolites and bacterial abundance acutely post-inflammation, we sought to measure whether they persisted into adulthood (P110). We first examined microbiome diversity as an indicator of overall lasting effect of DSS. At P110, the number of observed microbiota features remained significantly decreased in female DSS-treated mice compared to VEH (*p* = .0207) (Figure 6A). Additionally, although microbiota compositions (as calculated using Bray-Curtis distances) became more similar between DSS-treated and VEH mice, differences in sex and treatment remained significant (PERMANOVA, *p_sex_* = .001, *p_treatment_* = .004) (Figure 6B). Consistent with these long-lasting microbiota changes, analysis of colon histology revealed that DSS-treated colons at P110 retained several indicators of inflammation at sparsely found regions throughout the gut, specifically intestinal crypt hyperplasia and increased immune cell infiltration (representative images shown, Figure 6C). Nevertheless, the majority of the epithelium along the length of the colon appeared to have recovered after ∼40 days post cessation of DSS

**Figure 6.**
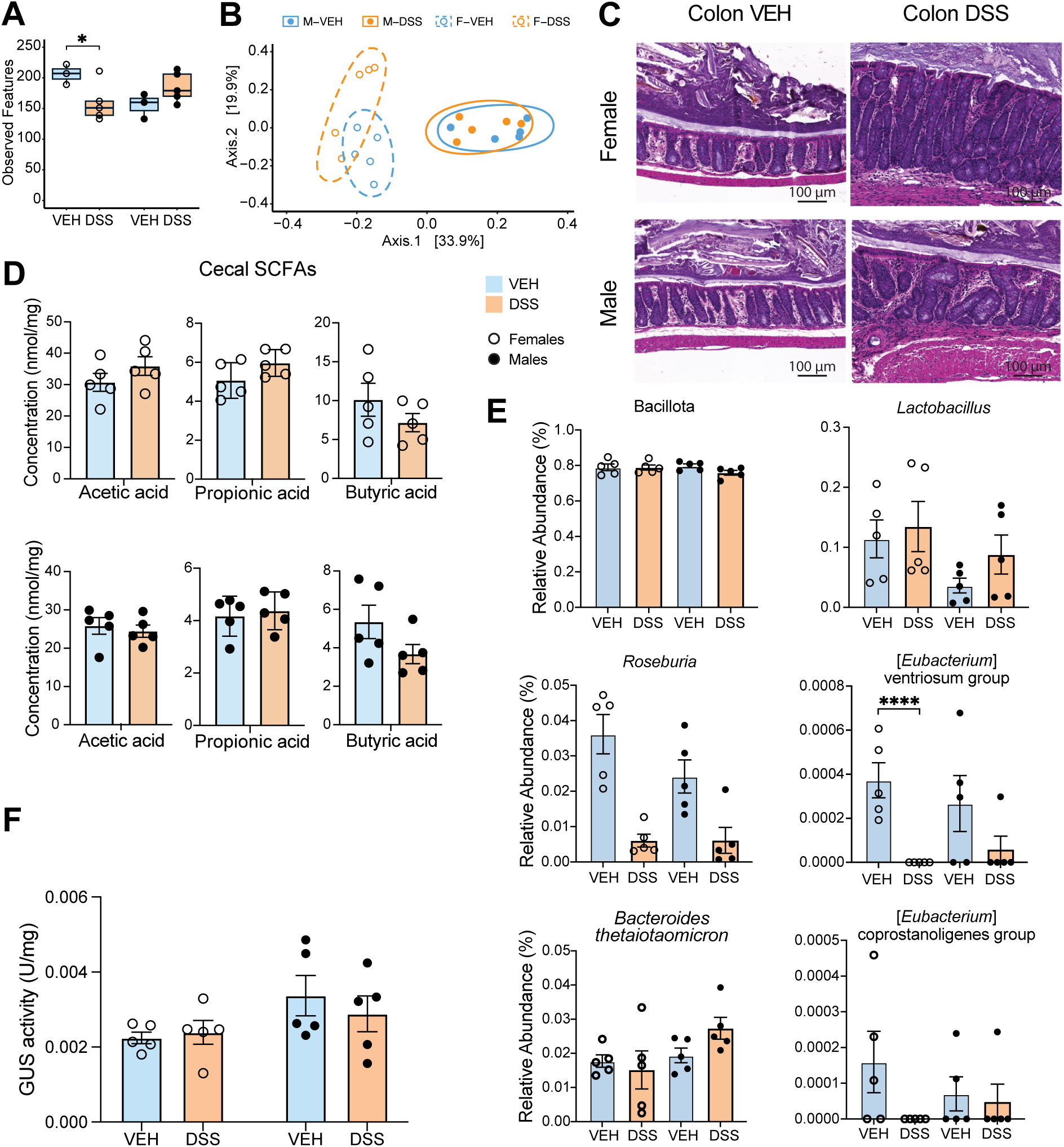
Early-life DSS treatment causes long-lasting physiological differences and changes to microbiota composition. *A.* Diversity of microbiota composition ∼30 days after cessation of DSS at P110, measured in number of unique features per sample. n=5/group. *B.* Principal Component Analysis of Bray-Curtis diversity index at P110. n=5/group. *C.* Representative histological images of hematoxylin and eosin-stained colons at P110. *D.* Cecal concentrations of SCFAs at P110. n=5/group. See Supplement for full set of cecal measurements. *E.* Relative abundance bar plots of select SCFA-producing bacteria. n=5/group. *F.* GUS activity in cecal contents at P110. Activity measured from cecal contents and normalized to total protein/sample. n=5/group. \**p*<.05, \*\*\*\**p*<.0001. Error bars in bar plots represent SEM.

Following these observations, we next sought to examine whether gut bacteria and microbial signaling molecules showed long-lasting changes after recovery. First, we found that there were no significant differences in cecal SCFA levels between VEH- and DSS-treated mice at P110 (Figure 6D). Consistent with these results, we found that many of the differentially abundant SCFA-producing taxa identified to be differentially present at P71 such as *Eubacterium* coprostanoligenes group and *Roseburia* were no longer significantly different between treatments. In addition, there were fewer significantly different taxa identified at P110 (Figure 6E) compared to the P71 (Supplementary Statistics Table, Supplemental Figure 3A) acute timepoint, indicating that the microbiome composition shifted towards recovery. Of the taxa that remained differentially abundant, *Eubacterium* ventriosum group (*p_females_* < .0001) is a known SCFA-producing taxa (Figure 6E). Second, we measured GUS activity from cecal contents at this timepoint and found that, consistent with a shift towards recovery, GUS activity was no longer depleted in DSS-treated animals (Figure 6F). Ultimately, these findings support the notion that early life inflammation causes acute changes to the microbiome and microbial signaling molecules, some of which may equilibrate in absence of further inflammation in adulthood.

### Early-life intestinal inflammation disrupts microglial morphology and brain connectivity in both sexes

SCFAs have been shown to alter microglial morphology and function (Erny et al., 2015). We therefore examined changes in microglial activation state in several brain regions. Using MicrogliaMorphology and MicrogliaMorphologyR (Kim et al., 2024a), we measured 27 morphological features from 12,318 microglia in the dorsal hippocampus (average of 513 per mouse) (Figure 7), 5,971 microglia in the hypothalamus (average of 426 per mouse) and 8,482 microglia in the cortex (average of 446 per mouse) (Supplemental Figure 4).We identified four clusters of different morphologies (Figure 7A, Supplemental Figure 4D) including microglia showing ameboid, ramified, rod-like and hypertrophic characteristics. Within the dorsal hippocampus there was a significant treatment by cluster interaction within each sex (Supplemental Table 1). Sidak corrected posthoc tests identified a significant decrease in ramified microglia in DSS-treated females (Figure 7B). In males there was a significant increase in hypertrophic microglia in the DSS-treated males. There were no significant differences in hypothalamic or cortical microglia between treatments within either sex (Supplemental Figure 4G, H). We further examined expression of cytokine genes and several immune receptors in brain tissue of DSS and VEH treated mice (Supplemental Figure 5). There were no significant differences between treatments within either sex, indicating that the sex-specific impacts of early life DSS treatment on microglia were not solely driven by neuroinflammation.

**Figure 7.**
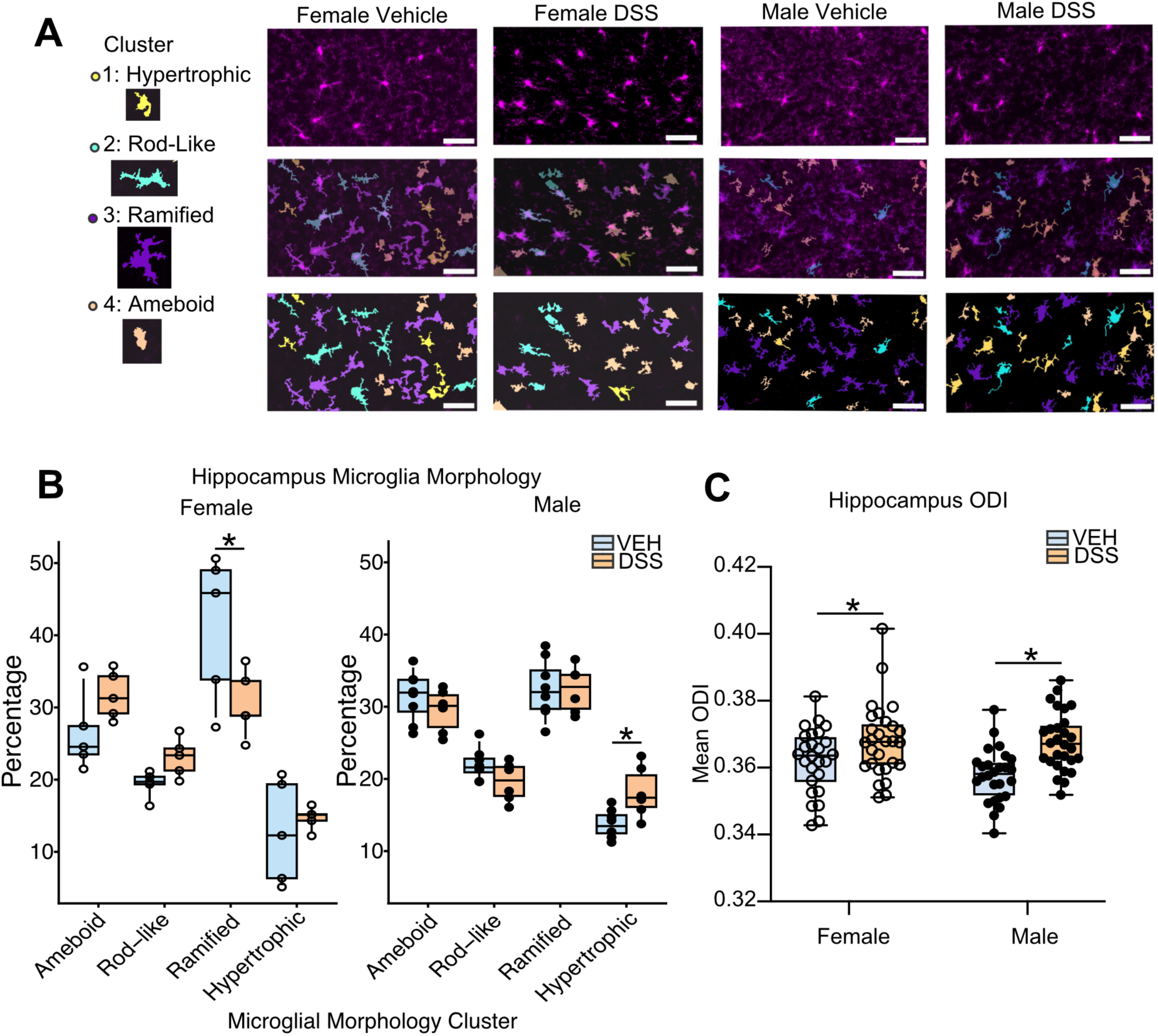
Long-Term Microglial Morphology and Extraneurite Microstructure Changes Following Early-Life DSS. *A*. Clusters were assigned to morphology descriptions based on relationship to individual metrics (see supplemental Figure 4). *B*. Example images from dorsal hippocampus of female (P110) and male (P72 and P110) vehicle and DSS treated mice reveals sex-specific changes in microglial morphology. Top panel is IBA1 staining. Middle panel is the overlay with individual cell morphology clusters and the bottom panel is the individual cell labels for each cluster. All scale bars are 50 µm. *B.* Quantification of the percentage of microglia within each cluster between treatment conditions for females and males. DSS significantly decreased the proportion of ramified microglia in females and increased hypertrophic microglia in males. n=4-8/group *C.* Orientation dispersion index (ODI) was calculated for the bilateral hippocampus across all samples. Data points are mean ODI values from each sample. Both males and females had significant increases in ODI in the DSS treatment when compared to treatment with the vehicle alone. n= 25-28/group * = *p* < .05, *** = *p* < .001.

### Clinical-translational quantitative neuroimaging reveals extraneuritic hippocampal microstructural changes following early life inflammation in both sexes

We performed ex-vivo multicompartment (MC) diffusion MRI and fit it to the neurite orientation and dispersion index (NODDI) model to assess and compare microstructural changes across biologically relevant regions (hippocampus, amygdala, striatum, thalamus, hypothalamus, frontal association cortex) between the VEH- and DSS-treated mice. MC diffusion MRI enables non-invasive characterization of neuronal cytoarchitecture at the mesoscale (1-100µm). Specifically, NODDI diffusion MRI permits non-invasive quantification of neuronal microstructure by measuring neurite density and cellular and morphometric changes occurring in the extra-neuronal space. In our imaging experiments, the mean orientation dispersion index (ODI, extra-neurite) and mean neurite density index (NDI, intra-neurite) values were tested using a 2×2 ANOVA considering the effects of treatment and sex. For ODI, a significant main effect of treatment was identified in the hippocampus (FDR-adjusted *p* = .0002, partial η^2^ = .1560) (Figure 7D). Post-hoc testing identified that both males and females had a significant increase in their ODI value when treated with DSS compared to VEH (males: Fisher’s LSD *p* = .0001; females: Fisher’s LSD *p* = .0269). Additionally, a significant main effect of treatment was found in the thalamus (FDR-adjusted *p* = .0095, partial η^2^ = .0822) (Supplemental Figure 6). Post-hoc testing indicated that females had a significant increase in their ODI value when treated with DSS compared to VEH, but that the increase in males did not reach significance (males: Fisher’s LSD *p* = .2353; females: Fisher’s LSD *p* = .0023). No significant treatment by sex interactions were identified in either the thalamus or hippocampus. No significant treatment or treatment by sex interaction effects were found in the frontal association cortex, amygdala, striatum, and hypothalamus. Furthermore, for NDI, no significant effects were identified in any of the regions of interest. Supplemental Table 1 contains the full findings for all regions tested.

### Early-life intestinal inflammation specifically during adolescence decreases seminal vesicle weight

To examine whether the developmental timing of one DSS treatment is sufficient to induce impairments in male sex organ development, we treated male mice with DSS for 5 days either at weaning (1xDSS Weaning) or during adolescence (1xDSS Adolescence) and collected seminal vesicles at P71 (Figure 8). We found that DSS treatment produced significant decreases in body weight (treatment x day interaction F(46, 318) = 15.111, *p*<.0001) following DSS treatment in both groups (Figure 8 and Supplemental Table S1 and S2). DSS treatment during adolescence significantly decreased seminal vesicle weights, highlighting this developmental period as specifically sensitive to inflammatory effects on endocrine function (F(2, 12) =7.514, *p*=.009, *p*=.0075). Conversely, DSS treatment during weaning did not alter seminal vesicle weight at P71 (*p*=.7565). Furthermore, there was no difference in testes weight between treatments F(2, 11) =.575, *p* =.5788.

**Figure 8.**
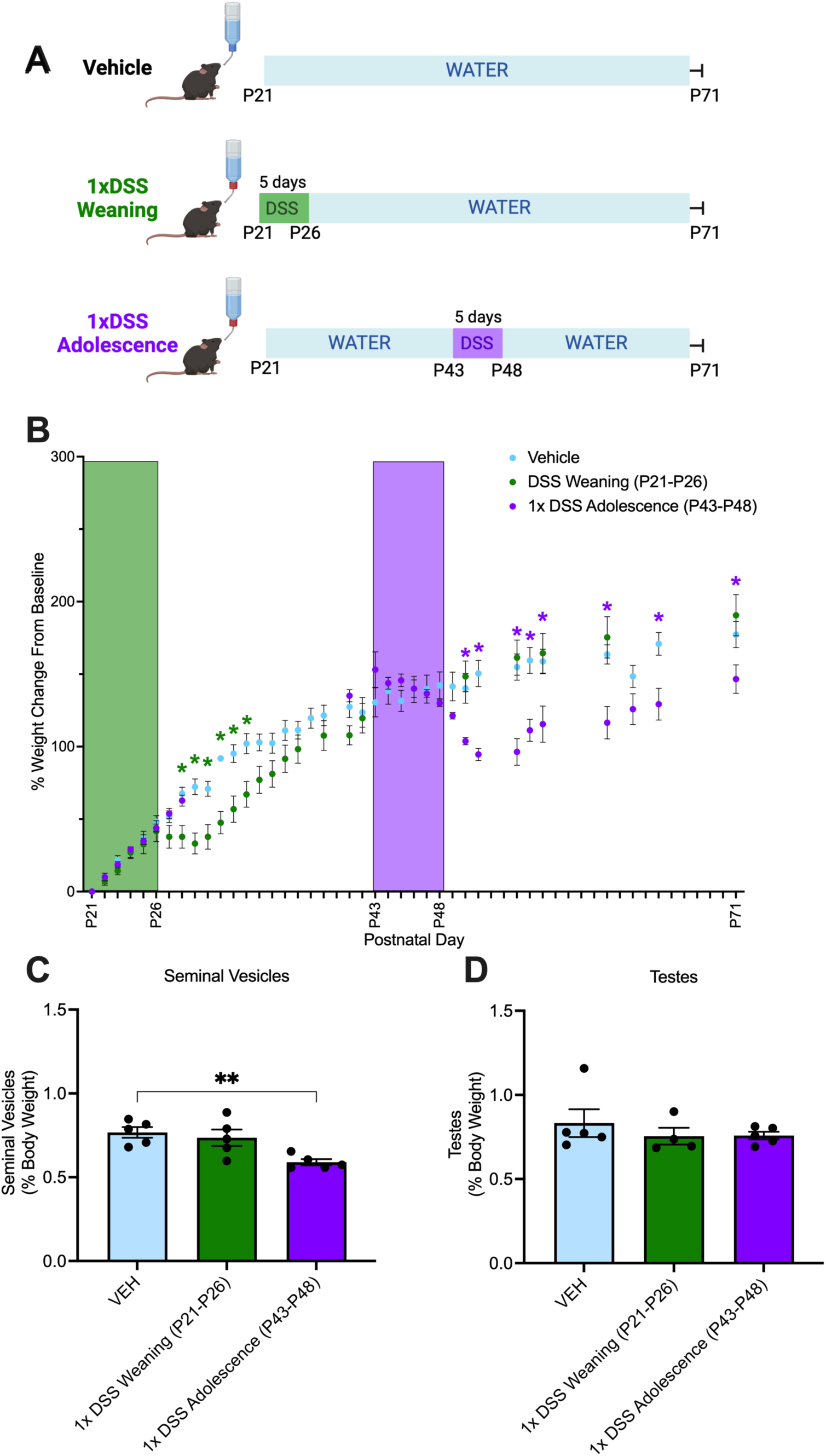
DSS Treatment During Adolescence Decreases Seminal Vesicle Weight. *A*. Male mice were treated with a single puse of DSS at either weaning (P21-26) or adolescence (P43-48) and were euthanized on P71. *B*. Body weight decreases following DSS treatment for the weaning group at P28-33 and following DSS treatment for the adolescent group beginning on P50 through the remainder of the experiment. *p<0.05 Sidak corrected posthoc tests. *C*. Seminal vesicle weights were significantly lower in the adolescent DSS treatment group compared to vehicle controls. *D*. Testes weights were not significantly different across treatments. n=4/group. Each graph bar and error bar represent the mean ± SEM. \**p*<.05.

## Discussion

The incidence of IBD is high and increasing, especially in early life (Benchimol et al., 2017, 2014; Ng et al., 2017)., and the disease is particularly challenging when it develops in children. Delay in the onset and progression through puberty is common in this population, and can result in reduced adult height and sexual immaturity (Gupta et al., 2020). In this study, we assessed the microbiome, intestinal, endocrine, brain, and behavioral effects of repeated DSS-induced gut inflammation in early life by developing a novel model of juvenile IBD. While early life gut inflammation did not impair the majority of the adult rodent behavioral repertoire, we identified a unique deficit in preference for female urine in DSS-treated male mice. These deficits in mate-seeking behavior corresponded to a decrease in seminal vesicle size, blunted circulating levels of androgen hormones and SCFAs, an altered gut microbiota, and altered hormone-modifying enzyme activity. Microglia were altered in the hippocampus, indicating hippocampal dysfunction may contribute to reproductive behavioral deficits in males (Kight and McCarthy, 2020).

In our model, DSS significantly decreased seminal vesicle weight (Figure 4A), circulating sex hormone levels (Figure 4D-E), and impaired male-specific preference for female urine (Figure 3C) – a key aspect of mate-seeking behavior in rodents (Keller et al., 2008; Malkesman et al., 2010). These differences were not driven by impaired olfactory discrimination, as both sexes and treatments performed normally on the olfactory habituation-dishabituation paradigm (Figure 3D-E). In both adolescent males and females with IBD, puberty onset is often delayed and progression through puberty is often prolonged, especially in patients with disease relapses (Hildebrand et al., 1994). Furthermore, adults with IBD report increased rates of sexual dysfunction (Bel et al., 2015), and infertility in both men (Hammami and Mahadevan, 2020) and women (Zhang et al., 2021; Zhao et al., 2019). This is consistent with findings from our study, where we found that DSS-treated male mice had decreased androgen hormones (Figure 4D-E) that may have directly contributed to the decrease in seminal vesicle weight (Yamane et al., 1986). Additionally, we observed that seminal vesicles in DSS-treated male mice were often not descended fully within the abdominal cavity (data not shown), and that a single DSS treatment during adolescence was sufficient to cause a decrease in seminal vesicle size (Figure 8C). While we did not observe direct impacts on male mounting behavior, future work will be required to fully examine the impacts of early life DSS on adult male reproduction and fertility.

Research suggests that hormone supplementation could potentially alleviate symptoms of IBD (Nasser et al., 2015; Rosen et al., 2015). However, the use of hormone supplementation in adolescents with IBD is contentious and can pose substantial risks due to potential side effects such as weight gain, mood changes, sleep disruption, and increased susceptibility to infections (Bishop et al., 2014). This necessitates new, non-hormone based therapies for treating adolescent IBD. The microbiota represents a potential lever for management of IBD-mediated alterations in hormones and development. Germ-free mice are known to have changes in sex hormones and pubertal onset (Markle et al., 2013; Weger et al., 2019), suggesting that the gut microbiome regulates sex hormone levels of the host. Furthermore, sex hormones can in turn regulate the gut microbiome and gastrointestinal transit time (Rastelli et al., 2022). This bidirectional relationship develops during puberty (Korpela et al., 2021). In both sexes, puberty is marked by increasing abundance of estrogen-metabolizing Clostridia and decreasing abundance of Bacteroidia (Korpela et al., 2021). Specific gut microbiota members can also secrete β-glucuronidase and β-glucosidase enzymes that deconjugate inactivated estrogens and androgens back to their active form (Ervin et al., 2019; Patel et al., 2023). These re-activated hormones are then secreted back into the circulating blood, where they can act throughout the body. In humans, estrogen levels are strongly and significantly associated with fecal Clostridia taxa, including three genera in the Ruminococcaceae family (Flores et al., 2012). This suggests that gut microbial families such as Ruminococcaceae may directly influence levels of host sex-hormones such as estrogen. Disruption of β-glucuronidase-producing microbiota results in a reduction in circulating estrogens (Baker et al., 2017), impairing the feedback loop between sex hormone levels and gut microbiota composition. In our novel model of pediatric IBD, we report differences in the key β-glucuronidase-expressing genus *Roseburia* acutely following the final round of DSS treatment, along with several *Eubacterium* groups that are known to deconjugate estrogens (Figure 5C) (Ervin et al., 2019). The relative abundance of these taxa were positively correlated with serum testosterone and androstenedione concentrations (Figure 5E), highlighting the potential involvement of these gut microbes in regulating systemic hormone levels. In addition, we observed significant decreases in cecal β-glucuronidase activity in DSS-treated males, with activity also trending lower in DSS-treated females (Figure 5G). Subsequently, we found that after resolution of inflammation, the microbiome shifts towards recovery – with most of the alterations in microbiome composition and microbial signalling molecules resolving (Figure 6). Together, these findings suggest that future microbiota-targeted therapeutics specifically impacting hormone-regulating microbes could potentially offset the negative effects of IBD on sexual health.

How changes in gut inflammation drive endocrine, brain and behavior changes is not well understood, but metabolite signalling molecules have been proposed as a potential link between all three systems (Silva et al., 2020). A number of metabolites have been implicated in IBD pathogenesis, including SCFAs, secondary bile acids (SBA) and tryptophan (Lavelle and Sokol, 2020; Sinha et al., 2020). These microbiota-derived metabolic signals can cross the blood-brain barrier (Mertens et al., 2017; Mitchell et al., 2011; Wenzel et al., 2020) and are critical for normal microglial development, gene regulation and metabolism (Erny et al., 2021, 2015; Thion et al., 2017). In our study, we found more differences in SCFA/MCFA abundances at the acute timepoint in the serum of DSS mice compared to VEH than in cecal contents (Figure 5A-B). This difference could be due to SCFAs being absorbed by host cells for use as energy sources. In particular, intestinal epithelial cells utilize SCFAs (particularly butyric acid) as a main source of energy, causing the concentration of SCFAs in the blood to decrease compared to what is measured in the cecum (den Besten et al., 2013). It has also been shown that butyrate uptake by colonocytes is impaired in inflamed IBD patient tissues (Thibault et al., 2010). Thus, DSS-induced inflammation could cause decoupling of cecal and serum SCFA concentrations. The reduced levels of some SCFAs observed in our pediatric IBD model may be responsible for the abnormal microglial morphology we observed (Figure 7B-C), with potential negative impacts on microglial regulation of neural circuit formation during formative years in childhood and adolescence (Sullivan and Ciernia, 2022). While previous work has shown that DSS can increase microglial inflammatory gene expression acutely in adult animals, with males showing a stronger response than females (Caetano-Silva et al., 2024), we did not observe changes in cytokine expression in our experiments (Supplemental Figure 5). However, prior work on the impacts of SCFAs on neuroinflammation is mixed. For example, increasing dietary fiber can offset the impacts of aging on microglial gene expression (Vailati-Riboni et al., 2022); however, a high fiber diet enhanced microglial inflammatory gene expression in an adult DSS model (Caetano-Silva et al., 2024) and acetate treatment exacerbated pathology in an Alzheimer’s disease mouse model (Erny et al., 2021), suggesting more work is needed to understand how SCFAs modulate microglia. We found decreases in valeric acid in both males and females treated with DSS. Valerate, a ester of valeric acid, suppressed inflammation induced microglial phagocytosis in an immortalized human microglia cell line (Wenzel et al., 2020). Consistent with these findings, we observed decreased proportions of ramified microglia in DSS-treated females (Figure 7B, indicating a decrease in homeostatic microglia. We also observed an increase in hypertrophic microglia in DSS-treated males (Figure 7B), a phenotype observed previously during neuroinflammation(Kim et al., 2024b). Together, these findings suggest that early life gut inflammation can alter metabolite signaling to the brain, influencing microglial morphology in a sex-specific manner. Future work will be needed to connect the alterations in SCFA directly to microglial morphological state shifts and functions related to behaviors during development.

In order to test the therapeutic efficacy of any future intervention in humans, diagnostic biomarkers and biomarkers of therapeutic response are critically needed. Herein, we demonstrated that early-life DSS treatment altered increased extra-neurite signal in NODDI within the hippocampus and thalamus, but not other brain regions. The NODDI model is has been used previously to calculate the diffusion signal specific to intra-neurite and extra-neurite compartments of the brain and is sentisitve to various neuropathological states (Barnett et al., 2019; Singh et al., 2023; Stowe et al., 2024; Yi et al., 2022). Consistent with our prior work (Yi et al., 2019), changes in the ODI signal alone are consistent with its sensitivity to microglial morphology and density (Figure 7C). Importantly, this imaging technique is being developed for use in humans (Garcia-Hernandez et al., 2022), paving the way for future studies to examine microglial changes in humans with IBD.

Finally, our main behavioral battery tested mouse anxiolytic, repetitive, cognitive, social and depressive-like behavior and found no significant differences between control and DSS-treated mice (Figure 2). This is in contrast to previous findings (Salvo et al., 2020), where 5 days of DSS at weaning (P21) resulted in adult (P56+) behavioral deficits on the novel object recognition task and light-dark task. This finding is rather unique in that the majority of adult DSS models show altered anxiety and memory only during active inflammation (Emge et al., 2016) and would be consistent with our findings that after recovery behavior is largely normal. One of the major differences between our work in Salvo et al., 2020 is that we used three repeated DSS treatments. The chronic nature of the current study may have induced resiliency and a faster recovery than in a single DSS treatment. Previous work in adult mice comparing one versus repeated rounds of DSS treatment found that acute-treated mice showed impairments in memory, repetitive, depression and anxiety like behaviors while chronically treated mice only showed deficits in repetitive behaviors (Matisz et al., 2020). The authors suggest that the lack of behavior deficits in the repeated DSS treatment group may have been due to an adaptive or tolerizing effect of repeated DSS cycles (Matisz et al., 2020). Similar findings in adult mice treated with repeated rounds of DSS found no changes in locomotion, anxiety, depression or compulsive like behaviors. However, they did find lasting alterations in social interactions (Brown et al., 2024), which we did not observe. Salvo et al. did not examine sex-specific behaviors in their model; in contrast, we have observed impacts specifically on male development. When we specifically treated adolescent mice with DSS we found impacts on seminal vesicle development (Figure 8C) similar to those observed after three rounds of DSS (Figure 4A), suggesting that DSS impacts on male sexual development are driven by impacts during adolescence. Future work is required to replicate and extend our findings to identify how timing of DSS treatment impacts reproductive health including the onset of puberty and life-long fertility.

The incidence of pediatic onset IBD disease has doubled in the last 10 years (Ashton and Beattie, 2024), and young children with IBD often have more severe disease and more frequent health visits than adults (Benchimol et al., 2011). Their illness also imposes a large emotional and financial toll on families and caregivers (Herzer et al., 2011). Developing effective treatments for adolescents with IBD holds great promise, as reducing inflammation early in the disease course may prevent long-term complications, need for surgery, and hospitalization. Our novel mouse model of pediatric onset IBD, which captures alterations in the microbiome, microbial signaling molecules and brain-behavior disruptsion, represents a unique avenue for advancing our mechanistic understanding of IBD in this key subpopulation. Our findings open the door for new treatment avenues for therapeutics targeted to adolescents with IBD, with minimal side effects and maximal benefits for growth, development, and quality of life.

## Acknowledgements

The authors acknowledge that the land we performed this research on is the traditional, ancestral, and unceded territory of the xwməθkwəy̓əm (Musqueam) Nation. The land it is situated on has always been a place of learning for the Musqueam people, who for millennia have passed on in their culture, history, and traditions from one generation to the next on this site. We encourage others to learn more about the native lands in which they live and work at https://native-land.ca/.

This work would not be possible without the help of numerous core facilities. In particular, the authors would like to acknowledge the staff at the Centre for Disease Modeling (CDM) at UBC, Ingrid Barta at UBC Animal Care Services (ACS) Histology, and the UBC Biofactorial High-Throughput Biology Facility, supported by the University of British Columbia Global Research Excellence Biological Resilience Initiative. Thank you to the Djavad Mowafaghian Centre for Brain Health NeuroImaging and NeuroComputation Core (NINC) for data storage and computational resources. This research was supported in part through computational resources and services provided by the Advanced Research Computing at the University of British Columbia. Additionally, we would like to thank the Kopp Lab at UBC for allowing use of their tissue processor, embedder, microtome, and slide scanner. Finally, this work received support from the BC Children’s Hospital Histology Core, Dr. Andy Sham, Dr. Catherine Chan, and the team from the Gut4Health Microbiome Core Facility at the BC Children’s Hospital Research Institute, Lisa Nikolai from the Chromatography Facility at the Faculty of Agricultural, Life & Environmental Sciences at the University of Alberta, and Dr. Ian Nation from Animal Pathology Services Ltd. (Canmore, Alberta). Thank you to Dr. Shelly McErlane for consultation on sex organ collection in mice. Thank you to the Centre for Disease Modeling staff and animal technicians for assistance with mouse experiments and animal care.

## CRediT authorship contribution statement

OS: Methodology, Formal Analysis, Investigation, Data Curation, Writing - Original Draft, Writing - Review & Editing, Visualization

CS: Methodology, Formal Analysis, Investigation, Writing - Original Draft, Writing - Review & Editing, Visualization

KN: Methodology, Formal Analysis, Investigation, Data Curation

SC: Investigation, Data Curation, Writing - Review & Editing

CR: Methodology, Formal Analysis, Investigation, Data Curation, Writing - Original Draft, Writing - Review & Editing

JEH: Investigation, Formal Analysis, Writing - Original Draft, Writing - Review & Editing

APS: Investigation, Formal Analysis

KL: Methodology, Investigation

JC: Investigation, Formal Analysis

JK: Investigation, Writing – Review & Editing

HY: Investigation, Formal Analysis

CAC: Investigation, Data Curation

NACG: Methodology

BDD: Investigation

NP: Formal Analysis

KV: Investigation

SA: Investigation

HG: Investigation, Writing – Review & Editing

TH: Formal Analysis

KS: Methodology, Writing - Review & Editing

JP: Resources, Writing – original draft, Writing– review & editing, Supervision, Project administration, Funding acquisition

CT: Conceptualization, Resources, Writing – original draft, Writing– review & editing, Supervision, Project administration, Funding acquisition.

AVC: Conceptualization, Resources, Visualization, Writing – original draft, Formal Analysis, Writing– review & editing, Supervision, Project administration, Funding acquisition.

---

**Supplemental Figure 1.**
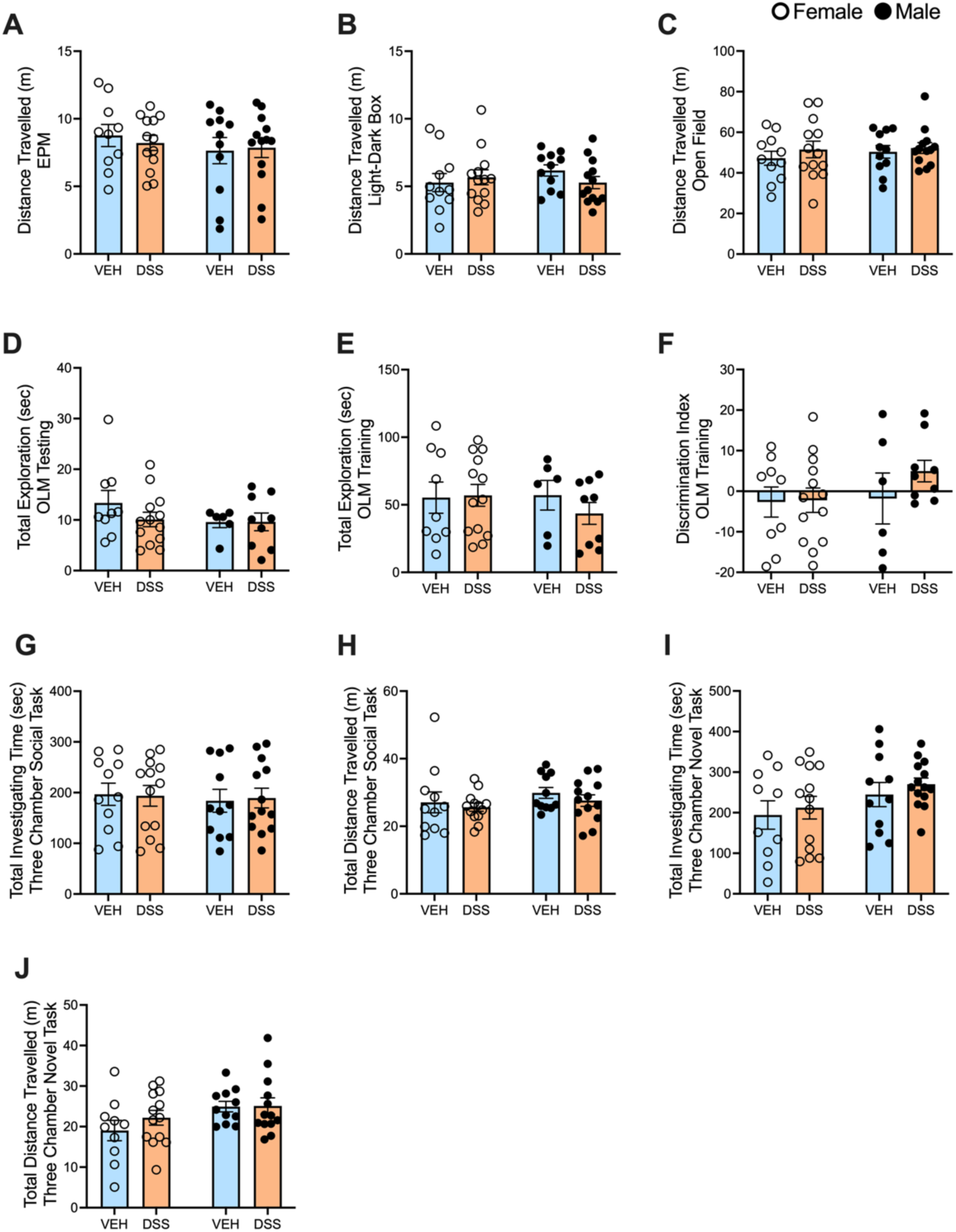
DSS treatment did not alter distance travelled or exploration behavior. There are no significant main effects of treatment or interactions with treatment and sex. *A*. Total distance traveled in meters (m) in the A. elevated plus maze n=10-13/group, *B*. light-dark box task n=11-13/group and *C*. open field n=11-13/group. Total investigation time in seconds for object location memory *D*. testing and n=6-13/group. *E*. training n=6-13/group. *F*. Discrimination Index for OLM training n=6-13/group. *G*. Total investigation time in the 3 chamber social task in seconds. n=11-13/group. *H*. Total distance travelled in meters for the three chamber social task. n=11-13/group. *I.* Total investigation time in the 3 chamber novel task in seconds. n=10-13/group. *H*. Total distance travelled in meters for the three chamber novel task. n=10-13/group. Each bar and error bar represent the mean ± SEM.

**Supplemental Figure 2.**
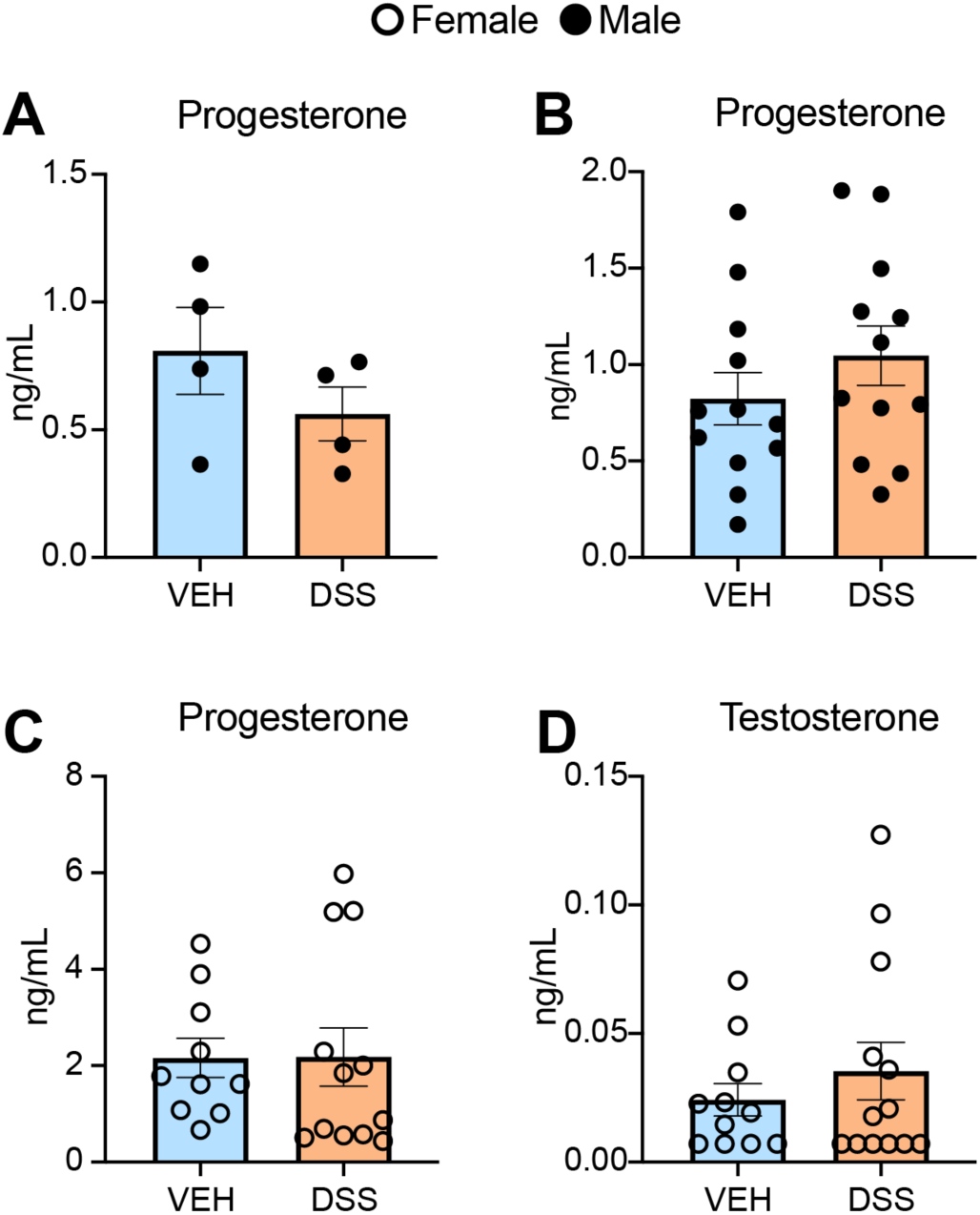
Effects of DSS treatment on male and female steroid levels. *A*. Three days after the last DSS treatment (P71), male serum progesterone levels were not significantly different between VEH and DSS treated males. n=4/group. *B*. Following the last behavioural test (P110), male serum progesterone levels were not significantly different between VEH and DSS treated males. n=12/group. *C-D*. Following the last behavioural test (P110), female serum progesterone (C) and testosterone (D) levels were not significantly different between VEH and DSS treated females. n=10-12/group. Androstenedione levels were non-detectable in VEH and DSS treated females. Each bar and error bar represent the mean ± SEM.

**Supplemental Figure 3.**
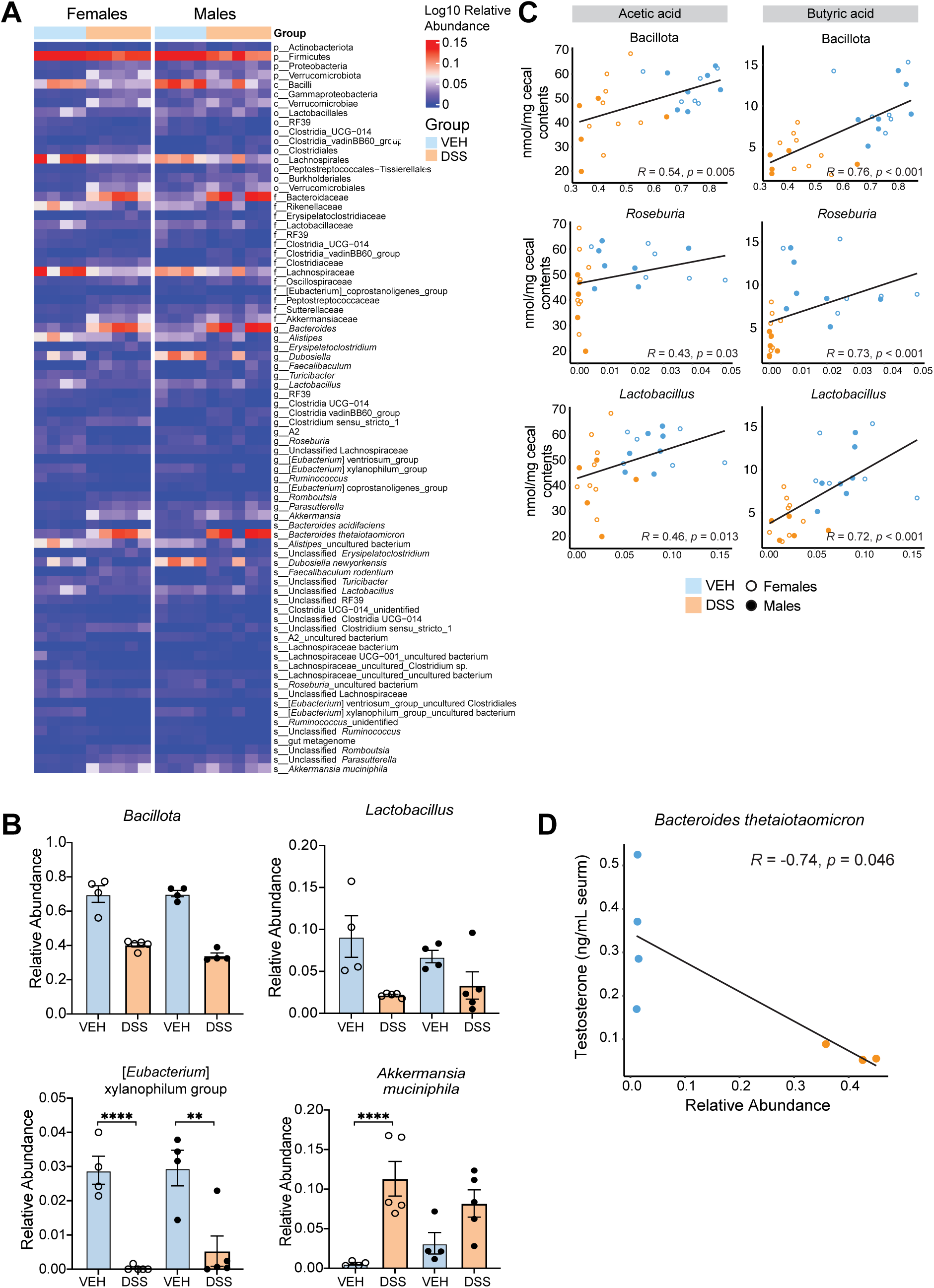
Relative abundance of select microbial taxa correlate with cecal SCFA and serum testosterone concentrations acutely following DSS. *A*. Heatmap of differentially abundant gut bacterial taxa identified by differential abundance testing at P71. n=4-5/group. *B.* Relative abundance of Bacillota, *Lactobacillus*, *Eubacterium* xylanophilum group, and *Akkermansia muciniphila*. n=4-5/group. *C.* Correlation of relative abundance of Bacillota, *Roseburia,* and *Lactobacillus* vs. acetic and butyric acid concentrations in cecal contents at P71. *R* values from Spearman’s correlation. n=5-8/group. *D*. Correlation of relative abundance of the species *Bacteroides thetaiotaomicron* vs. serum testosterone levels at P71. n=4/group. Error bars in bar plots represent SEM. \**p*<0.05, \*\**p*<0.01, \*\*\**p*<0.001, \*\*\*\**p*<0.0001.

**Supplemental Figure 4.**
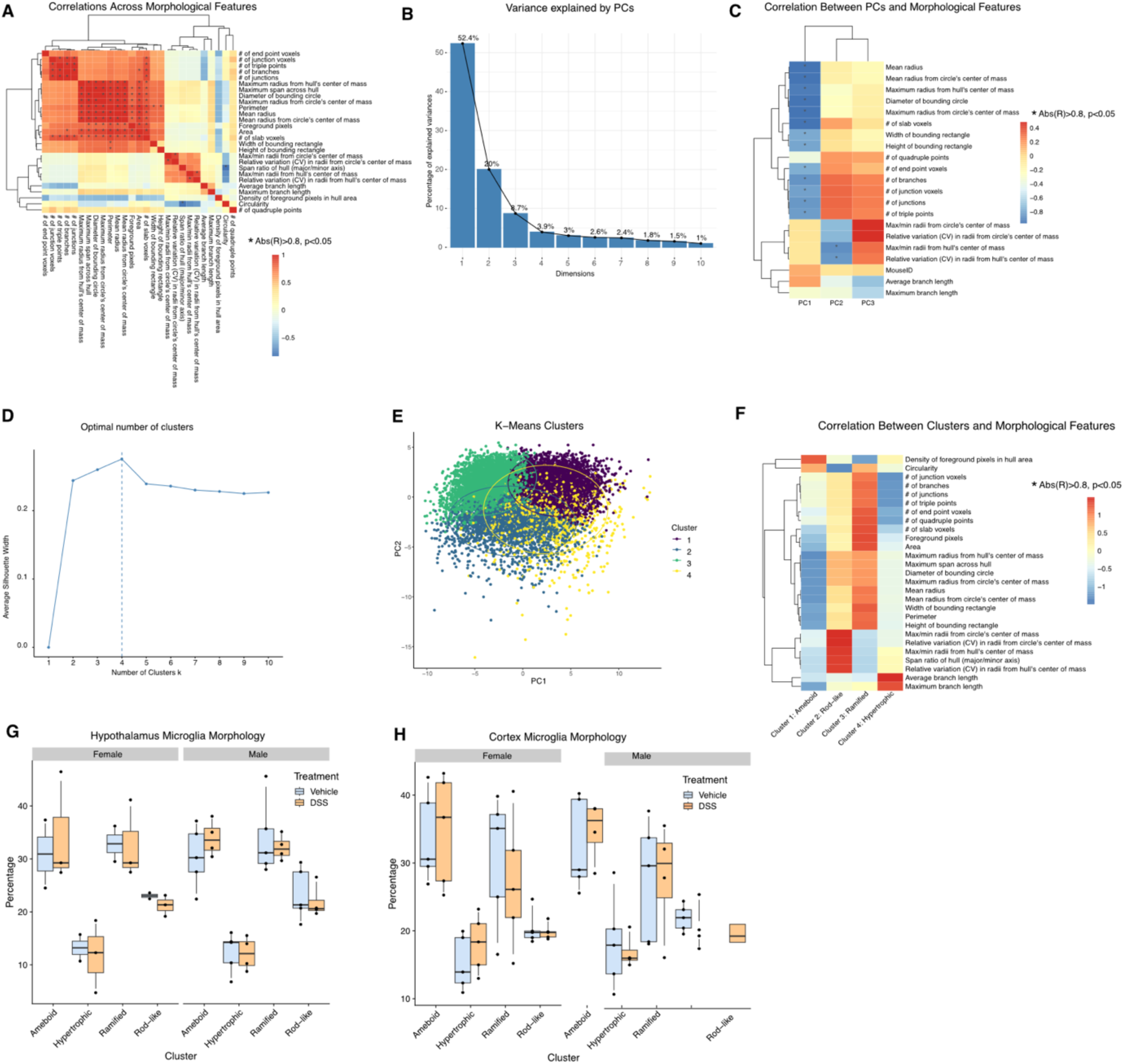
Microglial Morphology Analysis. A. Correlations between individual microglial morphology metrics. B. Percentage of variance explained by each principal component (PC) after clustering the 27 features. The first 3 PCs capture ∼85% of the total variance and were used for k-means clustering. C. The correlation between each PC and individual metrics. D. The silhouette plot reveals an optimal number of 4 clusters. E. K means clustering of 4 morphologyc clusters. F. Clusters were assigned morphology names based on their relationship to each individual metric. G. Percentage of hypothalamic microglia in each cluster between treatment groups for females and males. No significant differences were identified between treatments for either sex. n=2-5/group H. Percentage of cerebral cortical microglia in each cluster between treatment groups for females and males. No significant differences were identified between treatments for either sex. n=3-5/group.

**Supplemental Figure 5.**
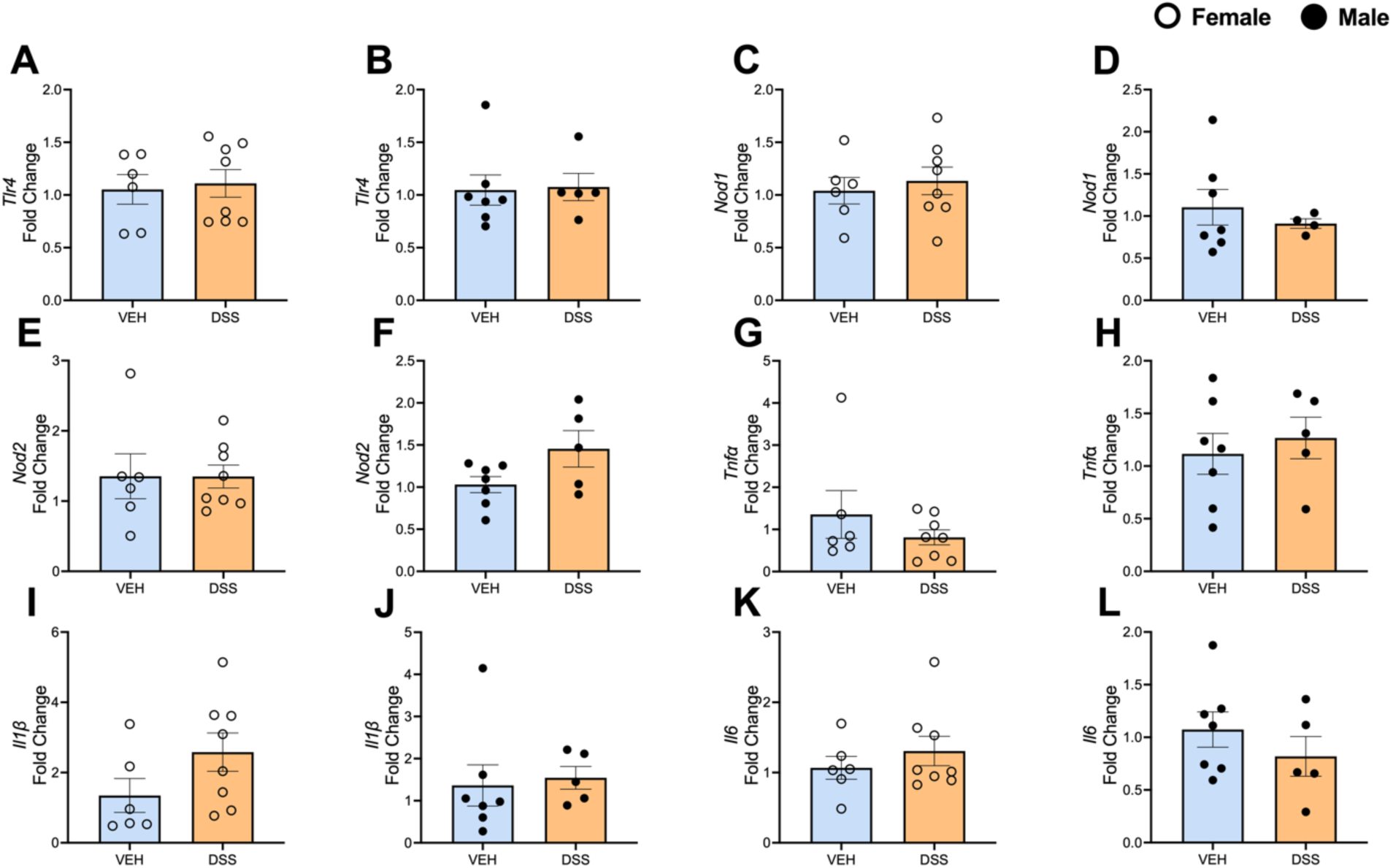
Gene Expression is Not Altered in Hippocampal Tissue from DSS and VEH treated male and female mice. Expression by RT-qPCR for *Tlr4* in A. females and B. males. Expression by RT-qPCR for *Nod1* in C. females and D. males. Expression by RT-qPCR for *Nod2* in E. females and F. males. Expression by RT-qPCR for *Tnfa* in G. females and H. males. Expression by RT-qPCR for *Il1b* in I. females and J. males. Expression by RT-qPCR for *Il6* in K. females and L. males. Fold change calculated to VEH controls within each sex for each gene. n=6-8/group. Each bar and error bar represent the mean ± SEM.

**Supplemental Figure 6.**
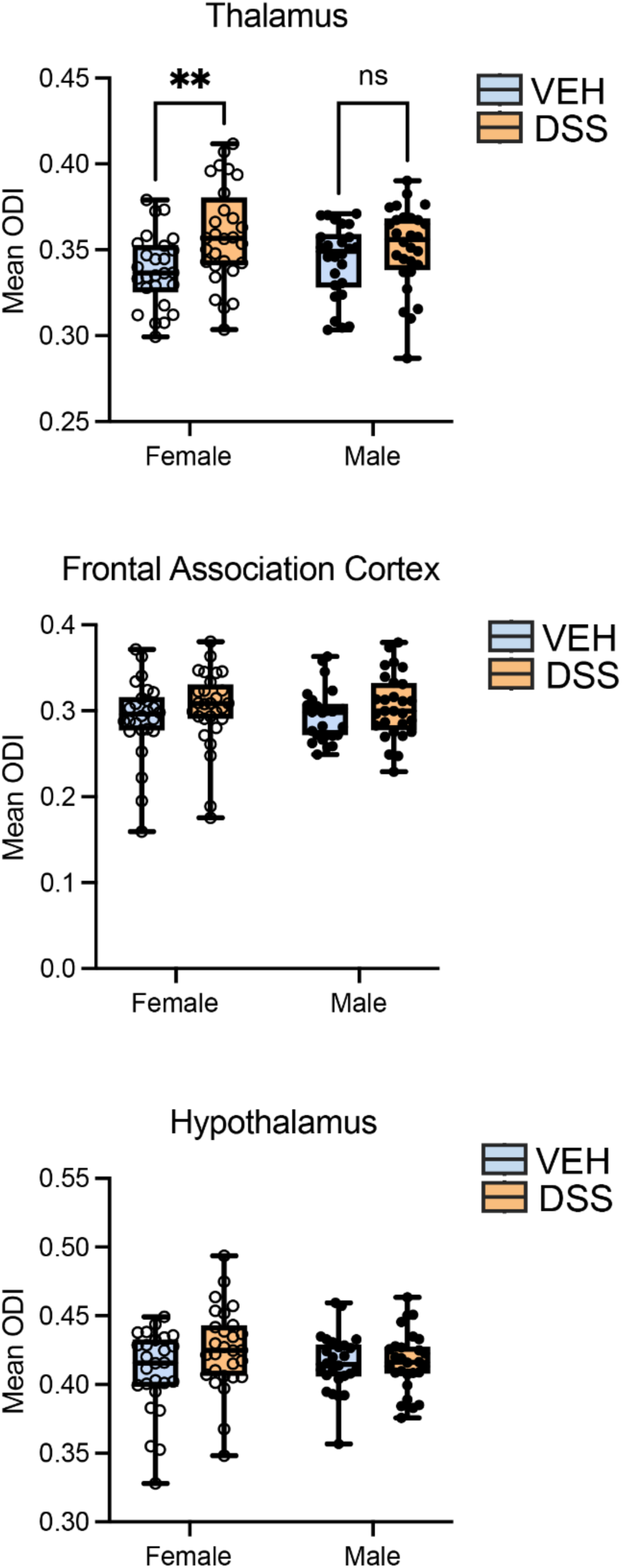
Orientation dispersion index (ODI) was calculated for the thalamus, frontal association cortex, and hypothalamus. Data points are mean ODI values from each sample. A significant effect of treatment was identified in the thalamus, and post-hoc tests found a significant increase for DSS females compared to VEH females. n= 25-28/group ** p < 0.01, Fisher’s LSD test.

